# Capacity and limitations of microfluidic flow to increase solute transport in three-dimensional cell cultures

**DOI:** 10.1101/2024.08.21.608799

**Authors:** Willy V. Bonneuil, Neeraj Katiyar, Maria Tenje, Shervin Bagheri

## Abstract

Culturing living cells in three-dimensional (3D) environments increases the biological relevance of laboratory experiments, but has the caveat of requiring solutes to overcome a diffusion barrier to reach the center of cellular constructs. We present a theoretical and numerical investigation that brings a mechanistic understanding of how microfluidic-culture conditions, including chamber size, inlet fluid velocity, and spatial confinement, affect solute distribution within 3D cellular constructs. Contact with the culture chamber reduces the maximally achievable construct radius by 15%. In practice, finite diffusion and convection kinetics in the microfluidic chamber further lower that limit. The benefits of external convection are greater if transport rates across diffusion-dominated areas are high. Those are omnipresent and include the diffusive boundary layer growing from the fluid-construct interface and regions near corners where fluid is recirculating. Less convection is required to approach an ideal maximally-supplied state when diffusion within the constructs is slow. Our results contribute to defining the conditions where complete solute transport into an avascular 3D cell construct is achievable and demonstrate how flow velocity must evolve with construct radius in order to maintain a given solute penetration depth.

## 1 Introduction

The morphology and behaviour of living cells fundamentally changes with the dimensionality of the environment they grow in. Three-dimensional (3D) cell cultures [1] more-accurately replicate the conditions that cancerous cells, migrating cells (e.g. , immune cells) [2, 3], or stem cells [4], amongst others, experience *in vivo*. Our understanding of organ development and tumour growth has been furthered by cell-culture techniques that promote the self-assembly of cells into spheroids or organoids (3D tissue constructs derived from stem cells) and subject those to controlled physico-chemical stimuli in order to modulate biological processes [5, 6]. These techniques also show promise to accelerate drug testing [7]. Here, we designate 3D tissue constructs grown in controlled environments as multi-cellular engineered living systems (M-CELS). M-CELS have a broader physiological relevance as they grow larger, because a larger size lets them mimic more-advanced development stages. Most M-CELS are engineered without a perfusable vasculature, which raises the problem of diffusion-limited growth. The incorporation of perfusable vasculatures into M-CELS is actively researched, but is a complex endeavour and lacks a generally applicable strategy [8, 9]. Nutrient transport and waste removal in avascular M-CELS rely on diffusion from their surface, which implies that cells beyond a certain depth do not receive enough nutrients and become necrotic i.e. , they prematurely die. Necrosis affects the phenotype of the remaining live cells, e.g. by releasing markers into the intercellular space that can induce inflammation or chemoattraction [10] and altering the mechanical properties of the live rim [11]. The balance between proliferating cells at the surface of M-CELS and dying cells at their centre confers a diffusion-limited maximum diameter to M-CELS [12]. Both phenotype alteration and size limitation restrict the stage of tumour growth or organ development up to which avascular M-CELS are a relevant biological model.

Microfluidic technology has enabled the culture of avascular M-CELS of larger diameters and greater compartmentalisation i.e. , of more-advanced development stages [13, 14, 15, 16]. The fluid velocities over M-CELS widely vary, as have the structures confining M-CELS, which are necessary for them not to be washed away. The magnitudes of the improvements brought about by microfluidic culture have been mixed. In one brain-organoid study, organoid diameters increased by a fifth under a steady fluid flow of 0.1 mm/s (our calculation, based on parameters listed in the paper), without necrotic-core prevention [13]. In another brain-organoid study, diameters increased by a third under oscillatory fluid flow of the order of 1 mm/s (our calculation), with necrotic-core prevention [16]. Fluid flow has been treated as an “on-off” parameter in all studies comparing static with microfluidic culture and its varying effects encourage the mechanistic elucidation of its effects on solute transport in M-CELS. Mathematical modelling and numerical experiments are well-suited to perform such an investigation because they allow to test and predic the performance of experimental setups in little time.

Previous modelling efforts on nutrient transport into M-CELS have yielded analytical expressions for nutrient concentration in systems with spherical [17, 18] or ellipsoidal [19] symmetry, isolated from their culture environment. Numerical work has been conducted on nutrient supply to spheroids in specific culture setups, including culture traps [20] or confinement pillars [21]. Some studies have applied optimisation algorithms to chamber geometries in order to maximise nutrient supply to M-CELS of a given cell type [22, 23]. Our systematic investigation of the effects of nutrient supply to M-CELS that is agnostic of cell type and chamber geometry will benefit the fundamental understanding of and the capacity to predict how chamber geometry and flow rate influence solute transport and necrosis occurrence in real M-CELS. Although nutrient supply is a primary interest, this investigation is applicable to any solutes, including drugs, that diffuse into an M-CELS and whose freely-diffusing form has a sink term: consumption or binding to an agonist receptor. We estimate not only the conditions for the prevention of necrosis, but more generally those for an increase in penetration distance. Our results highlight the penalising role of large diffusion-dominated areas such as corners or hydrogel capsules and of fast diffusion within M-CELS, which both delay the solute-transport enhancement brought by convection in the culture chamber.

## 2 Methods

### 2.1 Computational domains

Let a computational domain Ω comprise an M-CELS as an inclusion Ω_2_ in a straight fluid channel Ω_1_, such that Ω_1_ ∪ Ω_2_ = Ω. The interface between the domains is noted 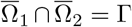. The fluid channel has an inlet ℐ and an outlet 𝒪. The channel walls were noted 𝒲_0_ at *y* = 0 and 𝒲_*h*_ at *y* = 2*h* and their union 𝒲. Ω_2_ was assumed porous and rigid with homogenous properties.

To isolate the effects of M-CELS confinement, we simulated two geometries. First, an “unconfined” geometry where Ω_2_ is a disc surrounded by fluid (Figure 1) and where Ω is bissected by a symmetry line S. The flow and the M-CELS are confined by the channel walls. The term “unconfined” refers to the M-CELS having no contact with any solid. Second, a confined geometry where Ω_2_ is a disc cut such that its height from the bottom of the channel was 90% of its diameter (Figure 2). Ω_2_ is in conctact with 𝒲_0_ through its linear boundary. This confinement is “geometrically minimal” in that it includes fluid-recirculation areas upstream and downstream of obstacles (in Ω_1_ close to the contact points Ω_1_ ∩ Ω_2_ ∩ 𝒲) and an area of contact between the M-CELS and a solid wall, but did not include wall-normal obstacles. The surface area of contact Ω_2_ ∩ 𝒲_0_ corresponds to the one observed in an experimental human lung-fibroblast spheroid (PB-CH-450-0811, PELOBiotech, Germany) cultured between confinement pillars in a microfluidic channel (Figure S2). In both geometries, Ω is described by a cartesian coordinate system (*x, y*) centered on the centre *O* of the smallest circle containing Ω_2_. A radial coordinate system (*r, θ*) is also defined from *O*.

**Figure 1.**
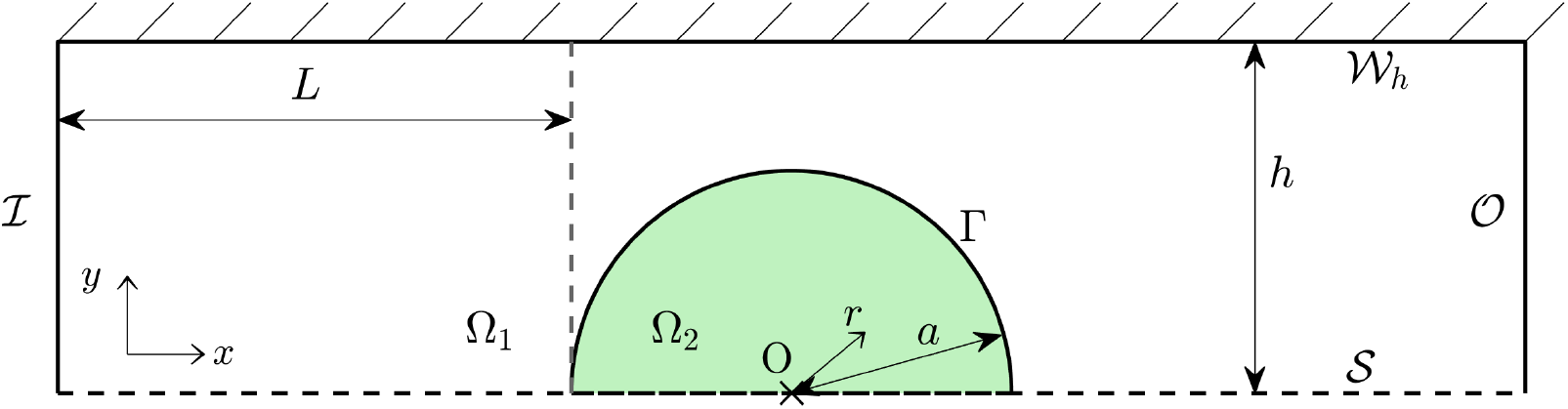
Computational domain (unconfined). M-CELS as a disc Ω_2_ of radius *a* in a rectangular channel Ω_1_. Ω bissected by a symmetry line 𝒮, parallel to the direction of flow. Coordinate origin at the centre of the M-CELS.

**Figure 2.**
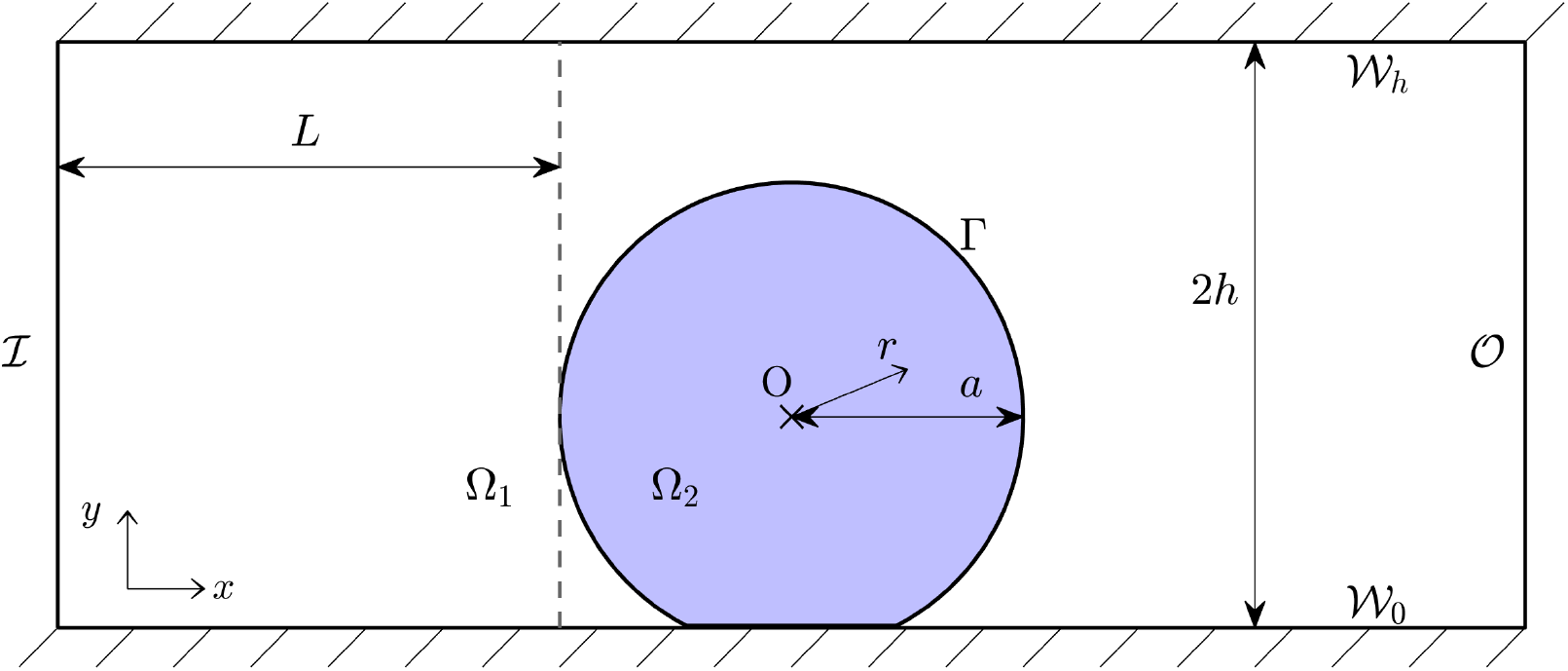
Computational domain (confined). M-CELS as a partial disc Ω_2_ cut such that its height above the bottom wall of the fluid channel is 90% of its diameter. Coordinate origin at the centr of the smallest circle containing Ω_2_.

In order to acquire an analytical understanding of the problem, we also define a quasi-1D “radial-linear” domain 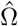 as the union of a radial section of a disc and the constant-width extension of that section (Section SM1).

### 2.2 Equations for fluid flow

Fluid flow in Ω_1_ is modelled by the steady Navier-Stokes equations with a parabolic inlet velocity and normal outlet flow (Supplementary Section SM2). The normalised momentum conservation equation is

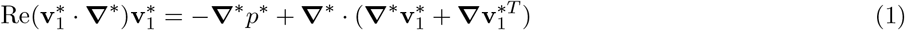

where the Reynolds number has been defined as Re = *ρv*_0_*a/µ*, local acceleration has been neglected, and non-dimensional variables are noted with asterisks. Microfluidic conditions involve peak inlet velocities on the order of 1 mm/s, M-CELS radii are at most of the order of 1 mm, cell culture medium has similar density and viscosity to those of water, so Re ≲ 1. We therefore did not neglect convective acceleration.

Fluid flow in Ω_2_ is modelled by the Darcy equations:

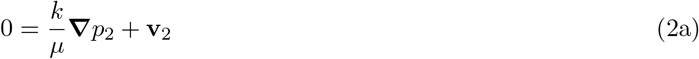

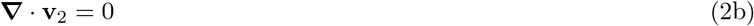

We assumed that the M-CELS had homogeneous properties, which is a valid assumption at least at early stages of M-CELS growth when little cellular differentiation has occurred. Continuity of pressure and velocity are prescribed at the fluid-porous interface:

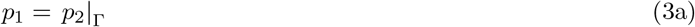

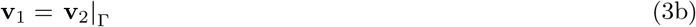

Pressure continuity may be applied at a fluid-porous interface when there is a separation of scales in the porous domain (here, the inter-cell distance of ∼ 10 nm is much smaller than the M-CELS radius of at least 100 µm[24]). The continuity of velocity between its Navier-Stokes and Darcy solutions results from mass conservation and the assumption that the slip length in Ω_2_ is zero. The latter scales with 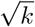 so is at most 100 nm and therefore much smaller than a cell (M-CELS are expected to be less permeable than the most permeable tissues, where *k* ∼ 10^−14^ m^2^ [25]).

### 2.3 Equations for nutrient transport

Solute transport was simulated by steady advection-diffusion-reaction equations. For *i* ∈ {1, 2},

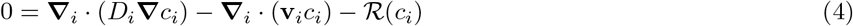

The velocities **v**_*i*_ are obtained from the fluid-flow equations. Diffusivities are uniform in each domain, at *D*_*i*_. Solute consumption is 0 in Ω_1_ and follows Michaelis-Menten kinetics in Ω_2_, as in models of tumour spheroids [26, 21]:

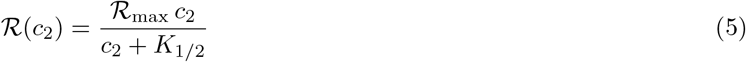

The consumption rate increases continuously from 0 in the absence of solute to ℛ_max_ at high solute concentration (*c*_2_ ⪢ *K*_1/2_). The half-rate constant *K*_1/2_ verifies ℛ (*K*_1/2_) = ℛ_max_/2. The consumption parameters are uniform in the M-CELS, which is reasonable at early stages of development. An unlimited solute supply, *c* = *c*_0_, was assumed at ℐ and 𝒪. This represents the common placement of a culture-medium reservoir on either side of a microfluidic chamber.

Let non-dimensionalised variables, noted with asterisks, be defined as 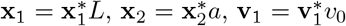, and 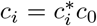 for *i* ∈ {1, 2}. To non-dimensionalise **v**_2_, we combine the pressure drop across a sphere in Stokes’ flow (Δ*p*_*S*_ ∼ *µv*_0_*/a*) and Darcy’s law (**v**_2_ ∼ *k/µ*Δ*p*_*S*_*/a*) to define 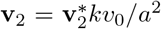. We also define 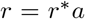 and 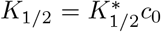. In Ω_1_, the non-dimensionalised transport equation is

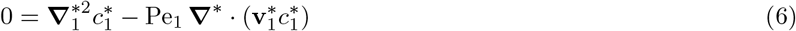

where the Péclet number is Pe_1_ = *v*_0_*L/D*_1_. In Ω_2_, the non-dimensionalised transport equation is

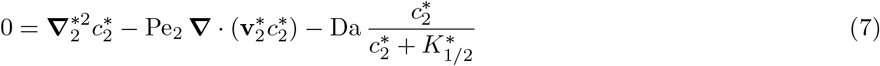

where the Péclet number is Pe_2_ = *kv*_0_/(*aD*_2_) and the Damköhler number is 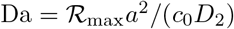.

The most favourable situation for convection in Ω_2_ shows the latter is negligible. Microfluidic flows rarely have a peak velocity higher than 1 mm/s, the permeability of M-CELS is assumed under 10^-14^ m^2^, and the question of solute transport sufficiency in M-CELS usually arises when those are larger than 100 µm. The Stokes-Einstein diffusivity of solutes of interest in M-CELS development is above 10^−11^ m^2^ s^−1^ (reached for proteins of 100 kDa) and we assume that the effective diffusivity in Ω_2_ for non-lipophilic solutes i.e. , those that do not cross cellular membranes, is at least 0.01 i.e. , *D*_2_ > 10^−13^ m^2^ s^−1^. This yields Pe_2_ < 0.1. Solute transport in Ω_2_ is described by a diffusion-reaction equation:

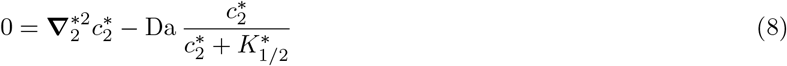

A no-flux condition is prescribed on the symmetry plane and walls:

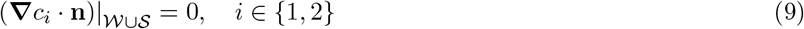

At the interface Γ, conservation of nutrient quantity implies continuity of concentration. Let the interface concentration be noted *γ*:

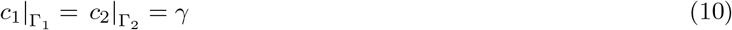

The evaluation of a quantity on Γ_*i*_ is understood as the limit of that quantity when approaching Γ from Ω_*i*_. Conservation of nutrient quantity also implies continuity of normal diffusive fluxes:

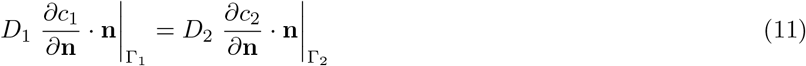

with **n** the normal to Γ (of arbitrary orientation).

### 2.4 Nutrient balance at the fluid–M-CELS interface

Three additional non-dimensional quantities are defined to describe physical balances of importance on Γ. The ratio of diffusive solute fluxes along the normal **n** (oriented into Ω_2_) is

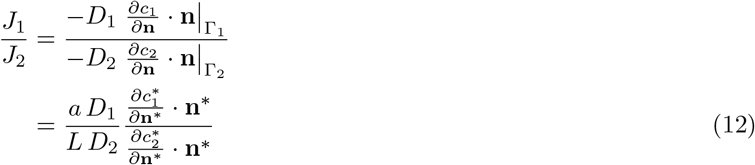

This ratio is equal to 1, so the ratio of interface concentration gradients is characterised by

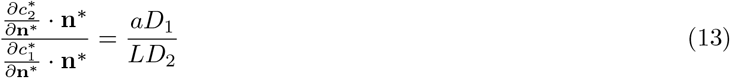

Let us define

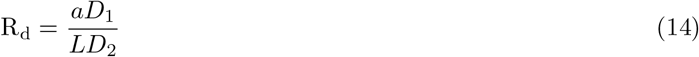

which characterises whether diffusion into Ω_2_ is supply-limited (low R_d_) or rate-limited (high R_d_).

Let us approximate *γ* as uniform to consider the solute quantity balance in a radial section of Ω_2_ of angle d*θ* centered on *O*. The corresponding section of Γ receives

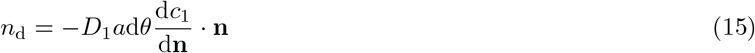

by diffusion, and

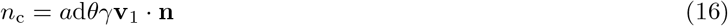

by convection. Let *r*_*n*_ be the radius where *c*_2_ falls to zero. In a radially symmetric geometry, Γ_*n*_ is the circle *r* = *r*_*n*_.

We allow *r*_*n*_ = 0 if *c*_2_ is positive everywhere. The radial section loses the following solute quantity due to consumption:

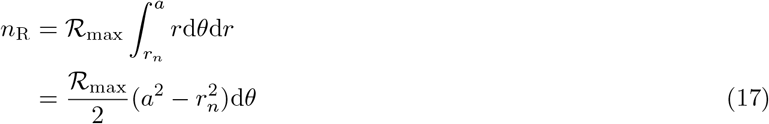

Let us now normalise the solute quantities and consider the ratio between diffusive supply to Γ and consumption in Ω_2_:

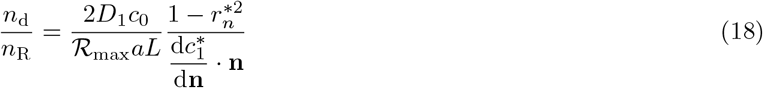

from which we define the diffusive supply number as

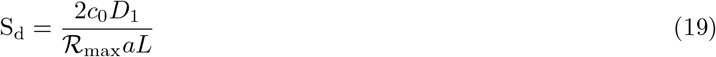

S_d_ describes how the diffusive supply of solutes to Γ affects the quantity of solutes consumed in Ω_2_. If S_d_ < 1, solute consumption is supply-limited i.e. , the quantity consumed by the M-CELS is limited by the quantity that it receives from its environment. If S_d_ > 1, solute consumption is rate-limited i.e. , the M-CELS consumes as much as its biology demands. S_d_ may be expressed as

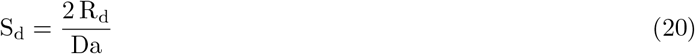

Let us consider the ratio between convective solute supply to Γ and consumption in Ω_2_:

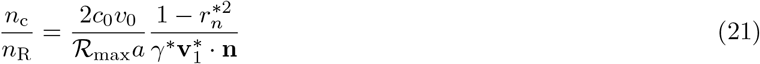

from which we define the convective supply number as

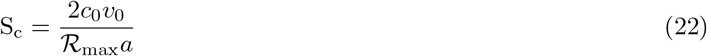

S_c_ may be described in an analogous way to S_d_ and expressed as

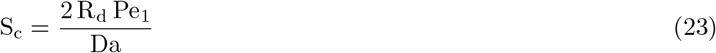

The ratios R_d_, S_d_, and S_c_ are measures of the excess or deficit of solute supply (diffusive or convective) relative to solute kinetics in the M-CELS (diffusive or consumptive).

### 2.5 Numerical methods

The simulations of coupled fluid flow and nutrient transport were implemented in the commercial software COMSOL Multiphysics^TM^ (COMSOL AB, Sweden, 6.1). The Supplmentary Section SM3 presents the details of that implementation and the dimensional parameter values.

### 2.6 Measures of solute transport

The goal of the simulations is to quantify the sufficiency of solute transport to cells in the M-CELS. The latter is measured by considering the shape and the surface area of the subset of Ω_2_ where *c*_2_ is sufficiently high. The boundary of sufficient transport (or necrotic boundary if the solute is a nutrient) is defined as:

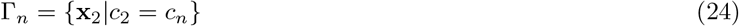

where *c*_*n*_ is a threshold concentration that depends on the metabolic needs of the cells. We use *c*_*n*_ = 10^−3^ *c*_0_ for generality. The center of mass of the zone of insufficient transport (or necrotic center if the solute is a nutrient) has the coordinates

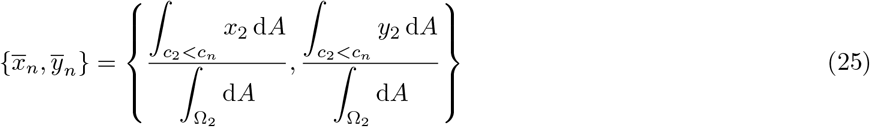

The fraction of sufficient transport is defined as the fraction of cells receiving solutes at a concentration above *c*_*n*_. It is termed the live fraction if the solute is a nutrient and is defined as

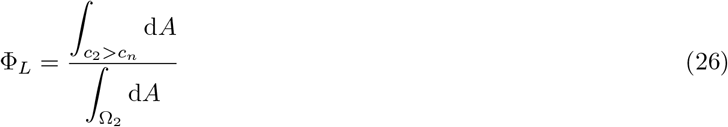

For simplicity and without loss of generality, the terms associated with necrosis are used in the following sections.

## 3 Results

The aim of microfluidic culture of rigid and avascular M-CELS, within which diffusion dominates solute transport, is to maximise solute supply to their surface. The concentration on an M-CELS surface should be as close as possible to its value in the reservoir, but this often is limited by the confinement of the M-CELS. Solute supply is reduced near M-CELS–wall contact areas where diffusive fluxes are restricted by constriction and fluid flow is recirculating or stagnant. This impedes the prevention of necrosis and limits the depth to which the penetration of a growth factor can be maintained. We first consider the effect of M-CELS confinement independently of solute supply i.e. , at *γ*^*^ = 1. This is termed the maximal-supply asymptote, where measures of solute transport are noted with an ∞ superscript. We then consider the effect of finite diffusive transport in static culture, termed the static asymptote. Measures of transport there are noted with a 0 superscript. We finally consider the role of convective supply i.e. , the trajectory that Γ_*n*_ and Φ_*L*_ follow between their static and maximal-supply asymptotes as convective supply increases. We distinguish supply insufficiency that occurs regardless of *γ*, which we term rate-induced, from insufficiency that occurs because *γ* is too low and that we term supply-induced. Maximising solute supply to the surface of M-CELS can either prevent supply-induced insufficiency, which is particularly interesting when the goal is to prevent necrosis, or reduce the zone of insufficient transport when the latter has a rate-induced component. That is not a satisfactory goal for nutrient supply, which needs to reach all cells, but is beneficial when the goal is a certain penetration distance for a non-nutritive solute.

### 3.1 Maximal-supply asymptote

At the radially-symmetric maximal-supply asymptote, the 𝒞^1^-regularity of *c*_2_ implies that *c*_2_ is zero with a zero gradient at the necrotic radius *r*_*n*_. This introduction of *r*_*n*_ has been done by others [17] and was shown to satisfactorily approximate Michaelis-Menten kinetics if 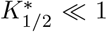 [26]. This is generally verified for cellular oxygen consumption. The solution of the transport equation (8) approximated with a constant consumption rate yields the following equation for 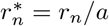 (details in Section SR1):

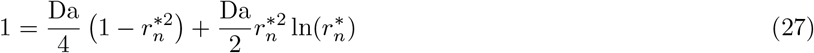

If 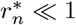, then 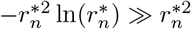. Equation (27) simplifies into a second-order polynomial of 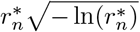. It admits a real solution if Da ≥ 4, which is positive and tends to 0 when Da → 4. Let Da_∞_ = 4 be the onset of rate-induced necrosis. The numerical simulations verified this in the unconfined geometry (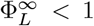 for Da > 4) and showed Da_∞_ = 3.4 in the confined geometry (Figure 3a). The penalisation of cell survival by confinement translates into e.g. , the maximum non-necrotic radius being 92% that of the equivalent unconfined M-CELS.

**Figure 3.**
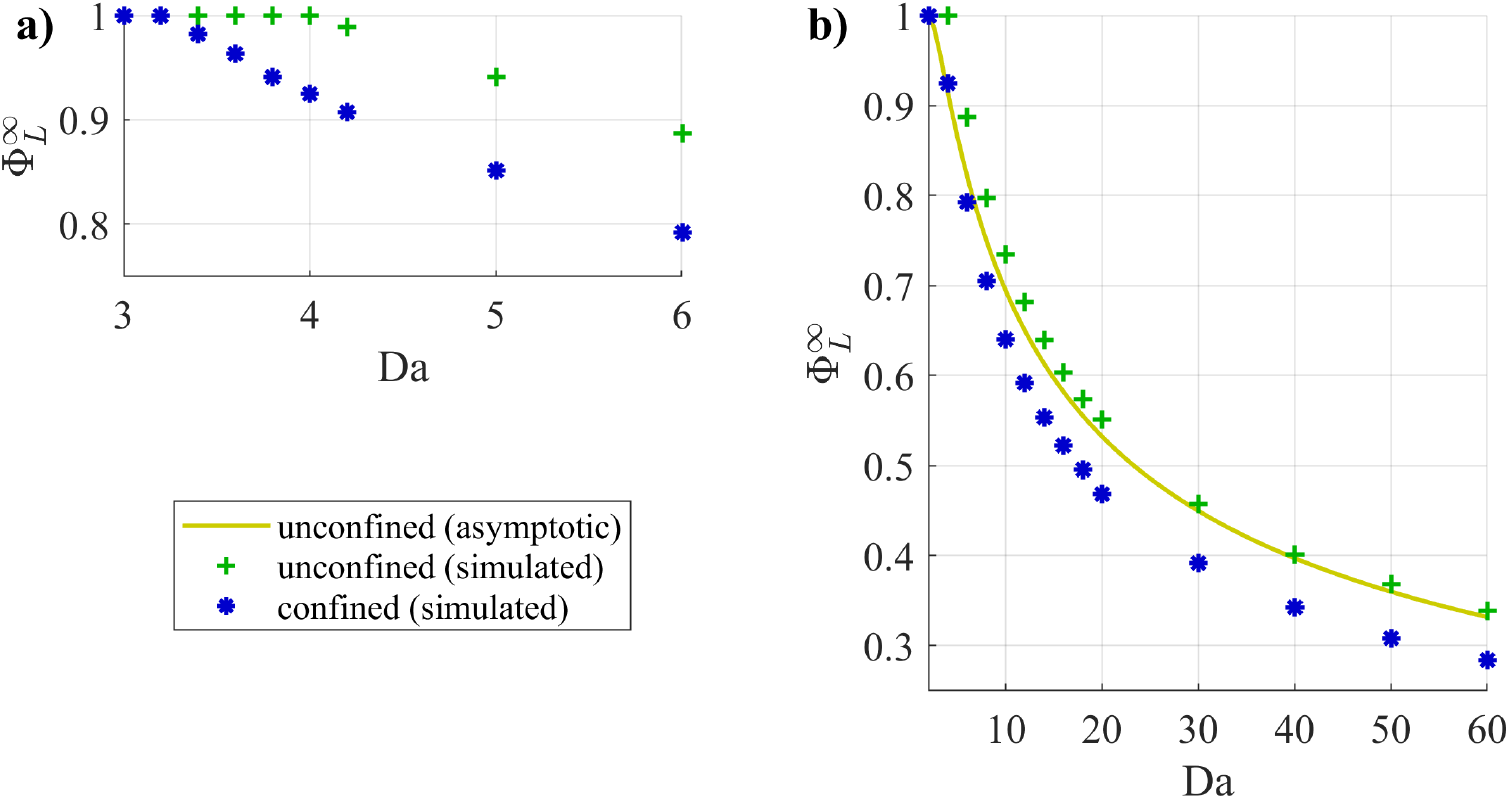
Live fraction at maximal supply: **a)** at the onset of necrosis, **b)** compared with its asymptotic expression at Da ⪡ 1.

If the necrotic area is large, 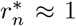 and Da ⪢ 1. Let 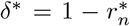. The logarithm term may be expanded around *δ*^*^ = 0 as

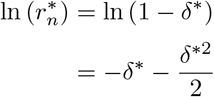

which yields the following expansion of Equation (27) at order 2 in *δ*^*^:

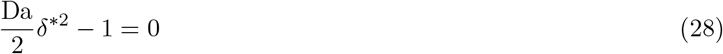

The asymptotic expressions for 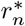 and 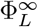 at high Da follow:

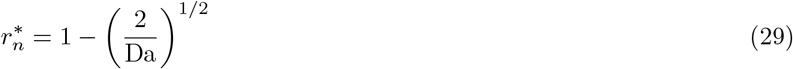

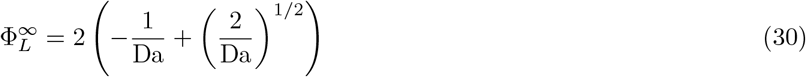

The latter is an excellent predictor of the unconfined live fraction for Da ≥ 10 (Figure 3b). The confined live fraction in the asymptotic high-Da regime was lower by around 15% for Da ≥ 10. This translates into necrotic radii being around 9% larger than in the equivalent unconfined M-CELS.

Figure 4 shows the shape of necrotic cores in confined M-CELS for varying Da. The contact surface skews the cores towards it. Those appear along the surface’s normal bissector when Da becomes larger than Da_∞_. Necrosis encompasses the M-CELS barycentre from its onset (see Γ_*n*_ at Da = 4). The normal to the contact surface passing by the barycentre thus is the region most vulnerable to necrosis.

**Figure 4.**
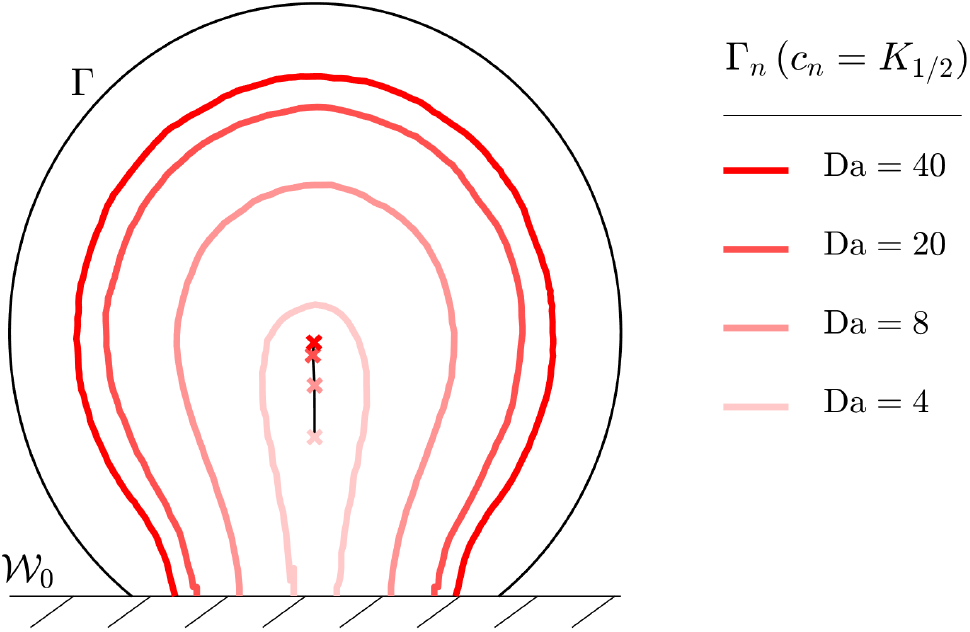
Necrotic boundaries at maximal supply in confined M-CELS and positions of the barycentres of the necrotic cores (marked by ×). Necrotic tissue in the area enclosed by Γ_*n*_ ∪ 𝒲.

### 3.2 Static asymptote

Let us obtain an analytical approximation of 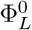 on the quasi-1D domain 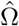. The non-dimensional necrotic radius 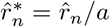 satisfies this equation:

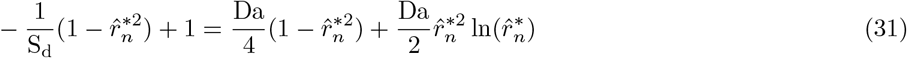

whose derivation is detailed in Section SR2.

#### 3.2.1 Prevention of necrosis

Let us investigate the prevention of supply-induced necrosis at Da < Da_∞_. When the necrotic core is small, 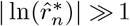, so (31) is approximated as

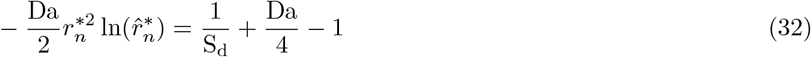

This admits a real solution if the right-hand term is positive. The necrotic core vanishes when that term becomes negative i.e. , with the unconfined Da_∞_ = 4,

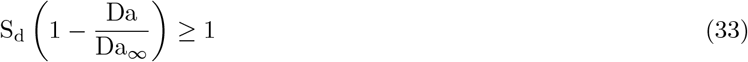

This satisfies physical intuition about extreme values of Da. When Da is low, it suffices that as many nutrients arrive at the interface as the M-CELS consumes (S_d_ > 1). As Da increases, *γ*^*^ needs to become ever closer to 1 to prevent supply-induced necrosis, which requires an excess of nutrient supply and S_d_ > 1. Finally, the value of S_d_ required for necrosis prevention diverges as Da approaches the onset of rate-induced necrosis. The condition (33) is expressed using the velocity scale of diffusion, which is *v*_dif_ = 2*D*/*d* at a diffusivity *D* across a domain of length *d*:

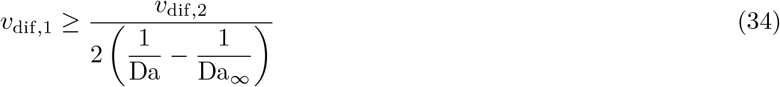

Necrosis prevention by large diffusive solute supply is counteracted by fast diffusion in the M-CELS. Preventing necrosis indeed requires raising concentration on the fluid–M-CELS interface and that concentration is lower when the diffusive-flux balance favours transport away from the interface, into the M-CELS. The diffusive-supply requirement is more stringent for small M-CELS with a fast metabolism than for large M-CELS with a slow metabolism. This applies to oxygen, whose diffusivity is on the order of 10^-9^ m^2^ s^−1^ in tissues but whose consumption rate varies over several orders of magnitudes between cell types ([18]).

The design constraints of experimental setups in practice lower the onset of rate-induced necrosis. They include space constraints on microscope stages and often room for the culture of multiple M-CELS. Those both impose a minimal value on *L*, of at least a few millimeters. Consequently, considering that *D*_2_*/D*_1_ is typically between 0.01 and 1, R_d_ is constrained under 10, and under 1 for nutrients like oxygen which readily cross cell membranes and for which *D*_2_*/D*_1_ ≈ 1. For oxygen, the maximal Da at which adequate nutrient supply can be achieved in static culture is Da ≈ 1, where the condition (33) is R_d_ > 2/3.

The condition (33) is verified by numerical simulations of the unconfined geometry (Figure 5a). Necrosis prevention is similarly achieved in the confined geometry, with S_d_(1−Da / Da_∞_) > 1 and Da_∞_ = 3.4 (Figure 5b). An approximately twice-higher diffusive flux to Γ is required to prevent necrosis relatively to the unconfined geometry. This corresponds to a reduction of the maximal permissible distance between a solute reservoir and an M-CELS by half in static culture, if all other parameters are equal.

**Figure 5.**
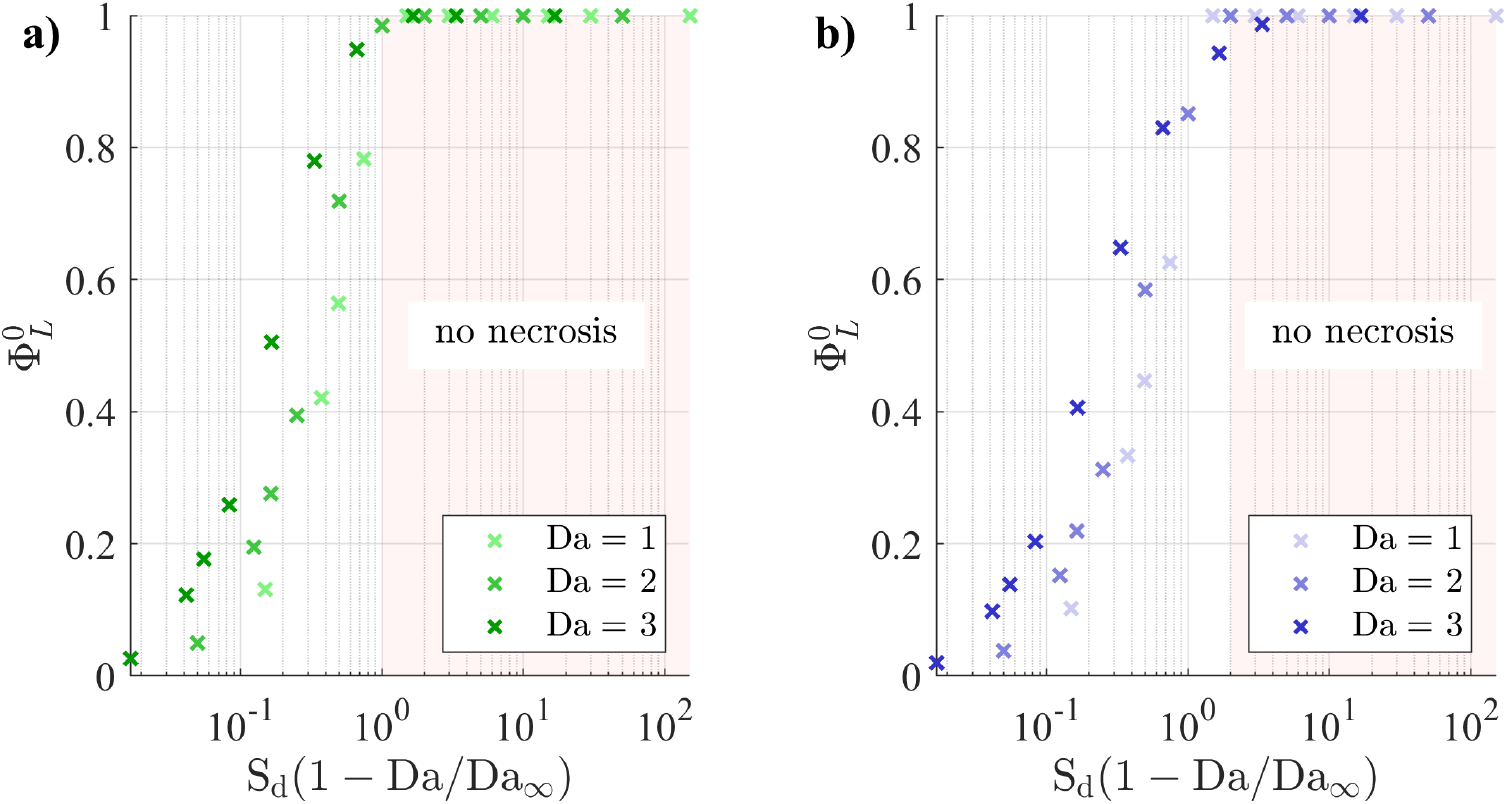
Live fraction at low Da and varying R_d_ in **a)** the unconfined geometry and **b)** the confined geometry.

#### 3.2.2 Approach of maximal-supply asymptote

Let us investigate the live fraction as a function of diffusive supply when there is rate-induced insufficient transport and diffusive supply is ample i.e. , S_d_ ⪢ 1, and R_d_ ⪢ 1. The live fraction normalised by its maximal-supply value is analytically derived in Section SR3 as

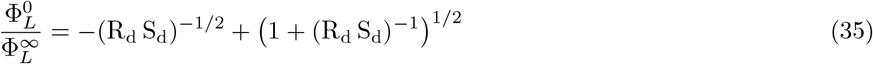

This describes a sigmoid function of S_d_ with an inflection point—the point where 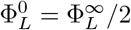 —shifted by R_d_. As for necrosis prevention, the excess of diffusive supply relative to consumption needs to be larger when diffusion into the M-CELS is supply-limited. The variable R_d_ S_d_ may be written

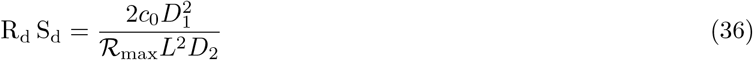

Counter-intuitively, the proximity of a static M-CELS with high Da to its maximal-supply asymptote does not depend on the M-CELS radius. It therefore does not depend on the M-CELS age if transport properties remain constant. The expression (35) also informs on how to adapt experimental parameters (*c*_0_ and *L*) based on those that may evolve with M-CELS development and health (ℛ_max_ and *D*_2_). For example, the tight alignment of neurons of the outer cell layer of cortical-brain organoids [27] increases *D*_2_ because the tortuosity becomes close to 1. If M-CELS diffusivity becomes 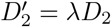, solute concentration needs to become 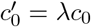 or the distance between the solute source and Ω_2_ needs to become 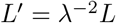 to maintain solute penetration.

Equation (35) is a good predictor of the simulated live fraction for Da ≥ 4 in both the unconfined and confined geometries (Figure 6). It overestimates the live fraction at low S_d_ because it does not account for the constriction between Γ and 𝒲_*h*_. Solutes at low S_d_ only reach the parts of Ω_2_ closest to I and O, so the uniform-*γ* assumption is less accurate. The live fraction progressed towards its maximal-supply asymptote along a similar trajectory in the confined geometry, but with a delay. S_d_ needed to be approximately 50% higher for the relative live fraction to progress to the same stage as if unconfined.

**Figure 6.**
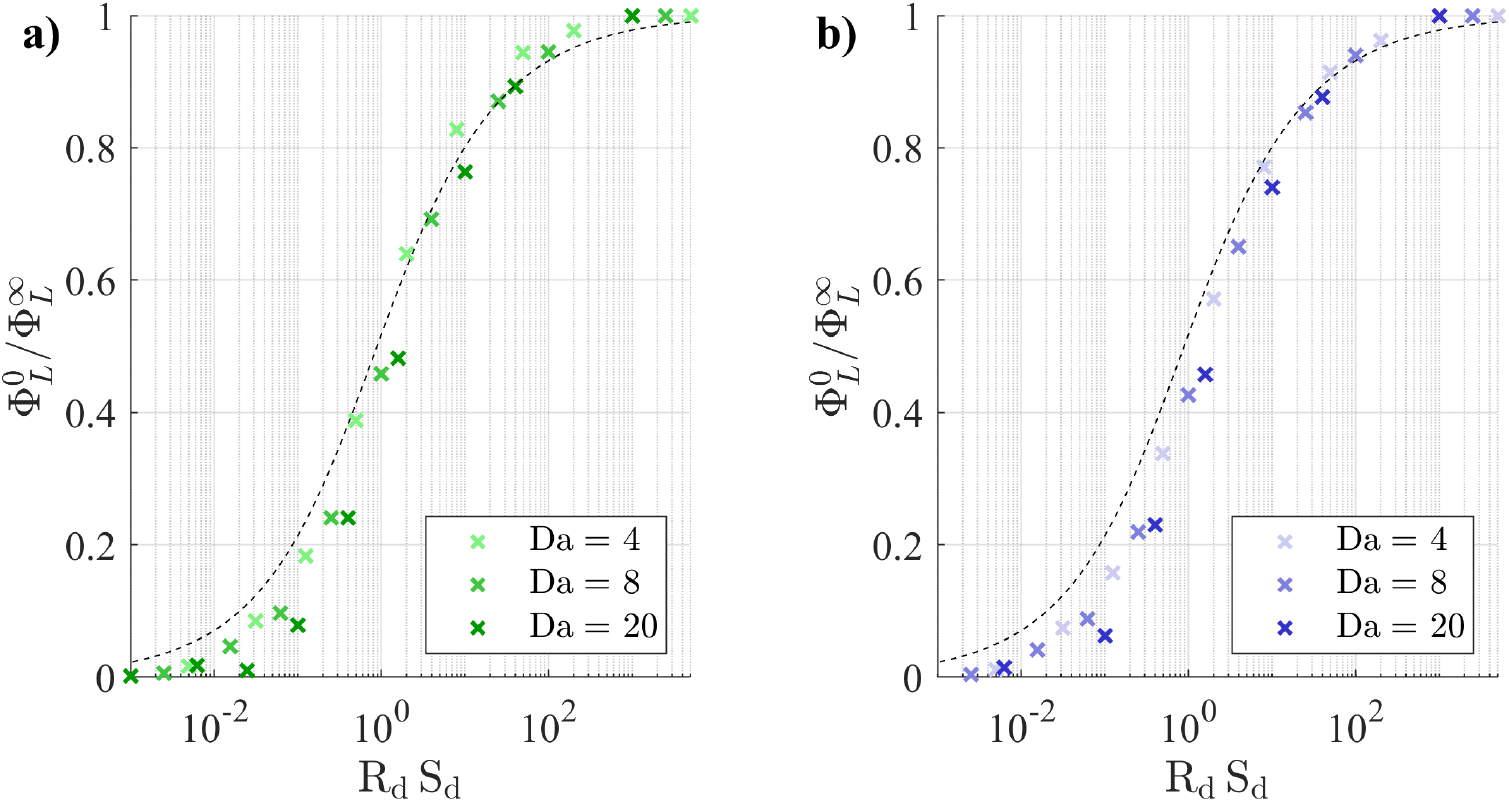
Live fraction at static asymptote for Da ≥ 2. **a)**. unconfined geometry. **b)**. confined. Dashed line: asymptotic expression (35).

At the static asymptote, the distance between the solute reservoir and the M-CELS implies that supply to the M-CELS surface is reduced. Preventing necrosis for Da close to the onset of rate-induced necrosis requires an excess of diffusive nutrient supply relative to nutrient consumption that is experimentally hard to achieve. The practical maximum Da in static culture can be lower than 1, which means a division by 2 of the maximum non-necrotic radius compared to maximally-supplied M-CELS. For M-CELS with rate-induced necrosis, the distance to the maximal-supply asymptote does not depend on their radius, so the efficacy of solute supply in a no-flow setup is constant for M-CELS with age-independent transport properties.

### 3.3 Microfluidic culture

#### 3.3.1 Function of increased convective supply

Convective solute supply only affects transport in the M-CELS if there is a deficiency at the static asymptote i.e. , S_d_ ≲ 1. Then, convection in Ω_1_ fulfills three distinct functions depending on Da (Figure 7). If Da ⪡ 1, concentration in Ω_2_ is approximately uniform. Any necrosis is supply-induced and can be prevented. It suffices that S_c_ > 1. If Da ⪢ 1, most of the M-CELS is affected by rate-induced necrosis. External convection would increase solute supply only to cells that are closest to Γ. The penetration distance of solutes would increase at most until its value at the unconfined maximal-supply asymptote 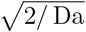, which still is poor transport. If Da ∼ 1, necrosis may be supply-induced or rate-induced, according to a threshold that depends on M-CELS confinement. Analogously to the static asymptote, supply-induced necrosis can be prevented by S_c_(1 − Da /4) ≳ 1. Rate-induced necrosis cannot be prevented, but may be reduced by high S_c_. External convection reduces the distance that solutes need to diffuse across to reach Γ, so it needs to increase when diffusive transport across areas of low fluid velocity is slow. Those areas are the diffusive boundary layer over Γ and the vicinity of the confinement corner (Γ ∩ Ω_2_). Since R_d_ S_d_ characterises the efficacy of diffusive supply, we postulate S_c_ ∼ (R_d_ S_d_)^−1^ when approaching the maximal-supply asymptote.

**Figure 7.**
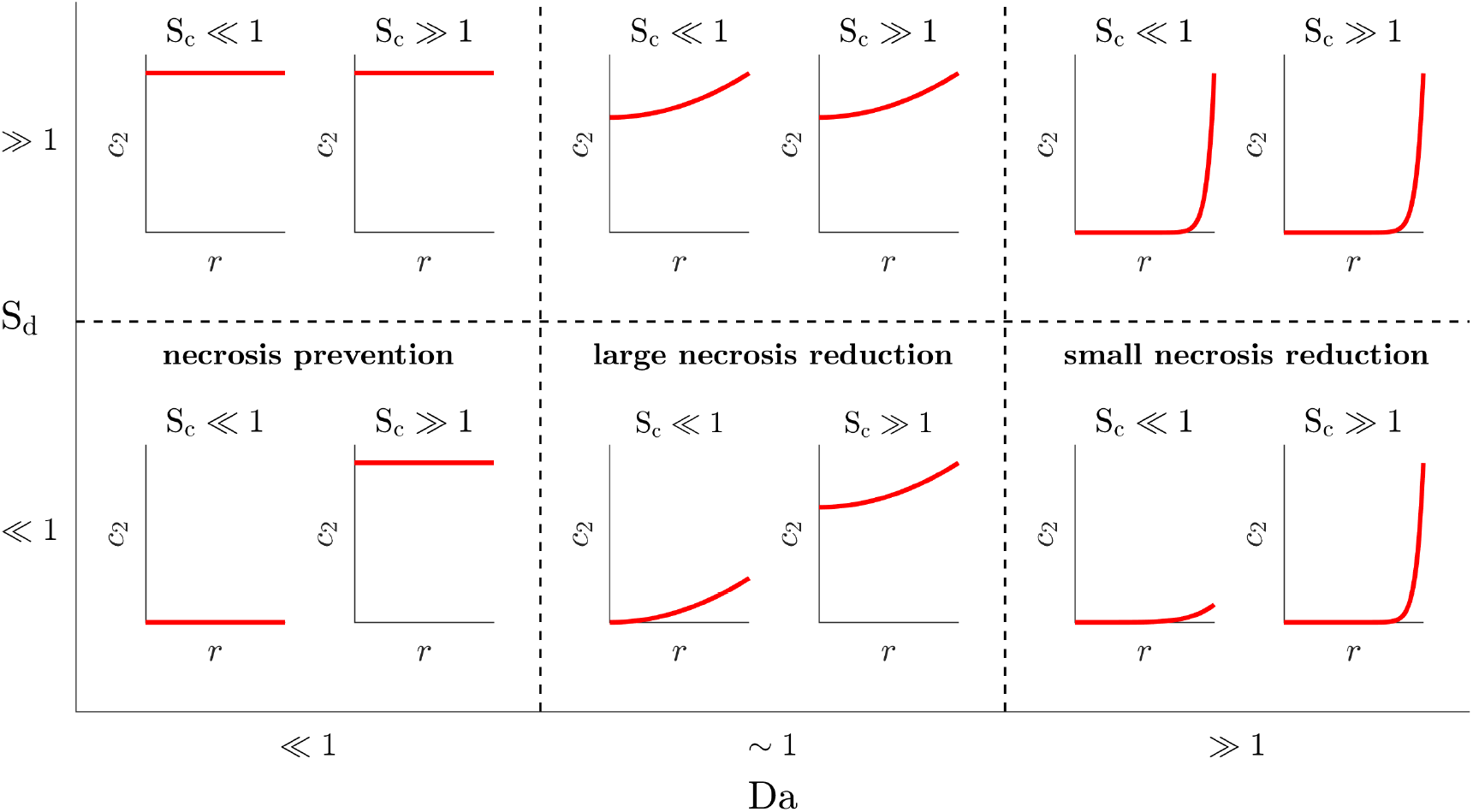
Qualitative effect of increased convective nutrient supply to Ω_2_ as a function of Da and S_d_.

#### 3.3.2 Evolution of solute supply with increasing convection

Let us describe how progressively-higher convection in Ω_1_ affects solute supply to Γ and its concentration in Ω_2_. At the static asymptote, solutes diffusing from ℐ and 𝒪 form a symmetric concentration field around Ω_2_ (Figure 8a). Low convection from ℐ to 𝒪 (Pe_1_ > 1 but S_c_ < 1) enhances solute transport from ℐ but inhibits it from 𝒪 (Figure 8b). The interface concentration *γ* increases on the upstream side of Γ but decreases on its downstream side. The area of insufficient supply, is shifted towards 𝒪. The supply increase on the upstream side of Γ mostly concerns its portion that is not affected by the confinement corner. Fluid flow near the latter indeed occurs as an infinite sequence of recirculation eddies of decreasing sizes and intensities [28]. Those were observed in the confined geometry because of the very large resistance to flow of Ω_2_, even though fluid transpires across Γ. Let Γ_*ρ*_ be the subset of Γ exposed to the recirculation eddies (Figure S5). Solute transport in that region is diffusion-dominated unless the convection in Ω_1_ is very large (the maximum velocity on the recirculation eddies decreases geometrically, with a ratio of ∼ 10^−3^ for the 60° angle of the confined geometry [28]). The efficacy of static solute supply thus remains determinant for the regions of Ω_2_ near the confinement corner, even when transport is convection-dominated in most of Ω_1_. As convection further increases (S_c_ > 1), solute flux from I becomes sufficient to fill the supply deficiency on the downstream side of Γ exposed to the primary flow and any remaining necrosis again becomes symmetric about *x* = 0 (Figure 8c). Supply increase to the confinement corner occurs as convection further increases and transport in the largest recirculation eddy becomes convection-dominated (Figure 8d).

**Figure 8.**
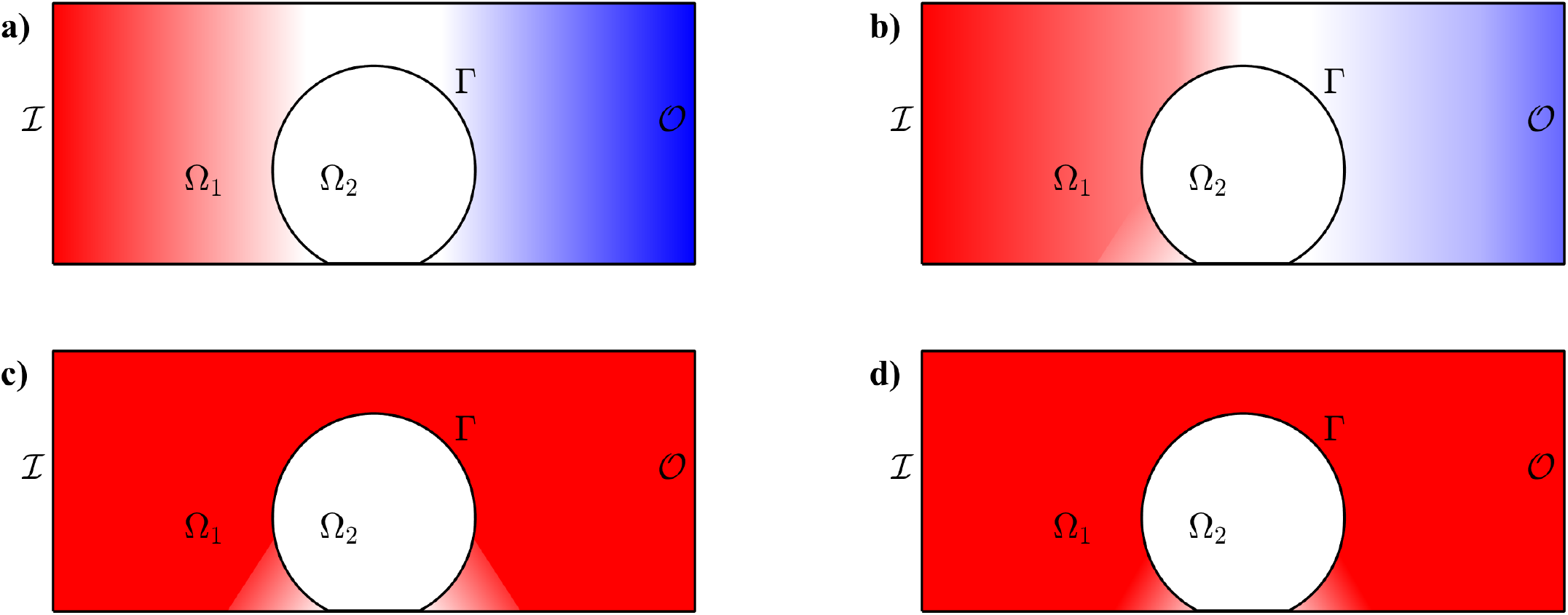
Schematic evolution of concentration in Ω_1_ under increasing convection from ℐ to 𝒪. Solute of origin on ℐ in red, on 𝒪 in blue. **a)**. Diffusive equilibrium with symmetric concentration around {*x* − 0}. **b)**. Low convection (S_c_ ∼ 1) increases concentration on the upstream side of Ω_2_ but decreases it on its downstream side. The upstream increase is limited near the confinement corner. **c)**. Convection of higher magnitude (S_c_ ⪢1) fills the primary flow region in Ω_1_ with solute (*c* ≈ 1 there), but not the confinement corners. **d)**. At ever higher convection, concentration increases in the confinement corner due to increased corner-eddy intensity.

#### 3.3.3 Prevention of necrosis

In the quasi-1D domain 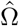, the non-dimensional necrotic radius 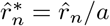 satisfies the following equation:

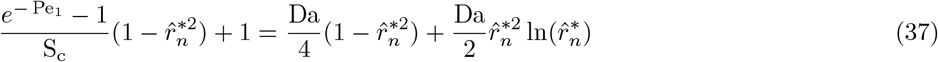

whose derivation is detailed in Section SR5. Let us assume that convection is sufficiently high that *c*_2_ is approximately symmetric and that Equation (37) may be used on Ω.

The necrosis-prevention condition on S_c_ is obtained from the same reasoning as for Eqaution (33) at the static-asymptote. Noting that 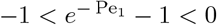, it suffices that

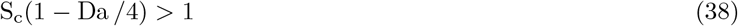

In terms of peak inlet fluid velocity, which is the most easily adjustable experimental parameter, this condition can be written

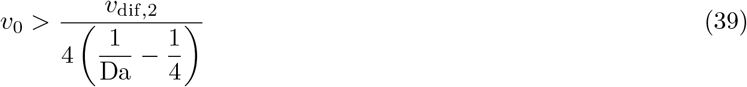

This depends on M-CELS properties only and not on diffusive transport in the microfluidic chamber. Like at the static asymptote, this condition expresses that necrosis prevention is penalised by fast diffusion in the M-CELS.

For an unconfined M-CELS of radius 0.5 mm where Da = 3 characterises oxygen kinetics, *v*_0_ ∼ 10^−5^ m/s. The corresponding inlet flow rate for typical dimensions of M-CELS culture chambers (1 cm × 1 mm cross-section, e.g. [15, 16]) is 1 ml/min, which is in the lower range of typical experimental flow rates. Flow rates are limited by the physiologic shear stress of the cultured cells, which is under 10^-3^ Pa at rest outside of blood vessels [29]. In order not to affect cell function in an M-CELS, peak microfluidic velocity thus should remain under 10^-3^ m/s (for a fluid viscosity of 1 mPa.s and a channel height of the order of 1 mm). That corresponds to 100 ml/min at typical channel dimensions. Necrosis prevention via external nutrient convection over an unconfined M-CELS therefore is achievable.

Numerical simulations showed that necrosis prevention was achieved if S_c_(1 − Da /4) > 5 in the unconfined geometry. (Figure 9a). The difference with the analytical condition (38) results from the diffusive boundary layer that grows from Γ into Ω_1_, which nutrients need to cross to reach Ω_2_. In the confined geometry, nutrients also need to diffuse across the corner recirculation areas, so a much higher excess of convection was required to prevent necrosis. That excess increased with lower R_d_ and higher Da, such that S_c_ ∼ 10^4^ was insufficient to prevent necrosis at Da = 3 (Figure 9b). This value is not advisable in experiments because it implies unphysiologic shear stress on cells. For oxygen in a confined M-CELS of radius 0.5 mm, the convection limit due to shear stress is at S_c_ ∼ 100. This means that necrosis prevention by microfluidic flow is possible if R_d_ ≥ 0.5 at Da = 2, but impossible if insufficient diffusive supply means that it exists at Da = 3.

**Figure 9.**
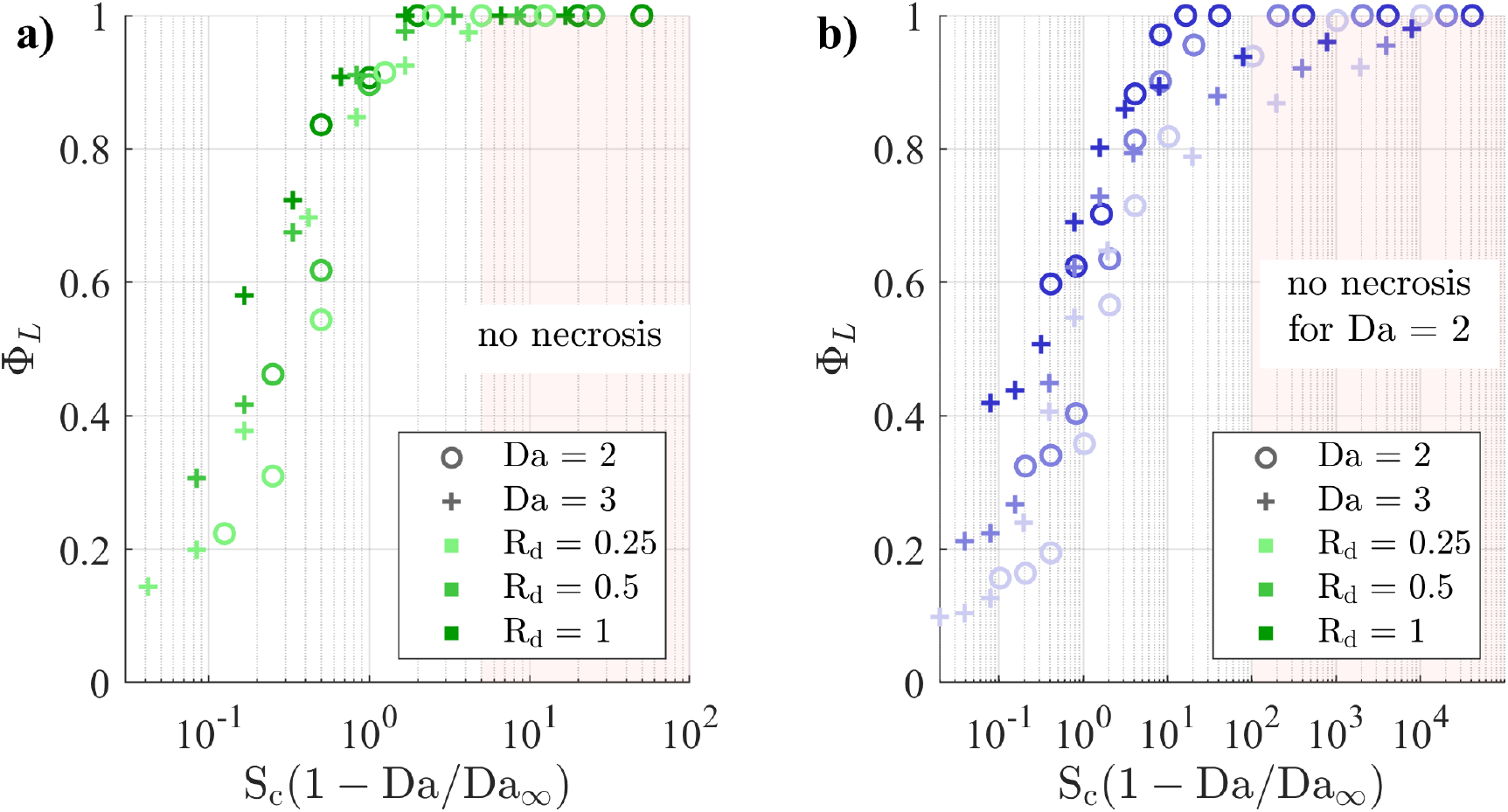
Simulated live fraction, normalised by its static and maximal-supply asymptotes, for Da such that any necrosis is supply-induced. **a)**. unconfined geometry. Necrosis prevention achieved for Da = 2 and Da = 3 if S_c_(1− Da /4) > 5. **b)**. confined geometry: necrosis prevention requires a larger excess of convective supply when R_d_ is low and Da is high. It is not achieved in any of the simulated cases when Da = 3.

#### 3.3.4 Transport asymmetry under moderately high convection

The asymmetry of solute supply to Γ when Pe_1_ > 1 and S_c_ < 1 translates into asymmetric necrotic cores, as the example of Da = 5 and R_d_ = 0.5 shows (Figure 10). The shape of the cores follows the progressive supply increase to Γ as convection in Ω_1_ increases. The cores first shrink on the upstream half of Ω_2_ (Γ∩{*x* < 0}), before shrinking along *y* as the supply deficiency is filled in the constriction between Γ and 𝒲_*h*_, and finally shrinking on the downstream half of Ω_2_. In the confined geometry, the third phase is followed by a reduction of the cores near the confinement corners. In that phase, ever-increasing convection in the primary flow region increases the intensity of the first recirculation eddy and thereby convective solute fluxes near the parts of Γ exposed to the recirculation region (Figure 10b and 10d).

**Figure 10.**
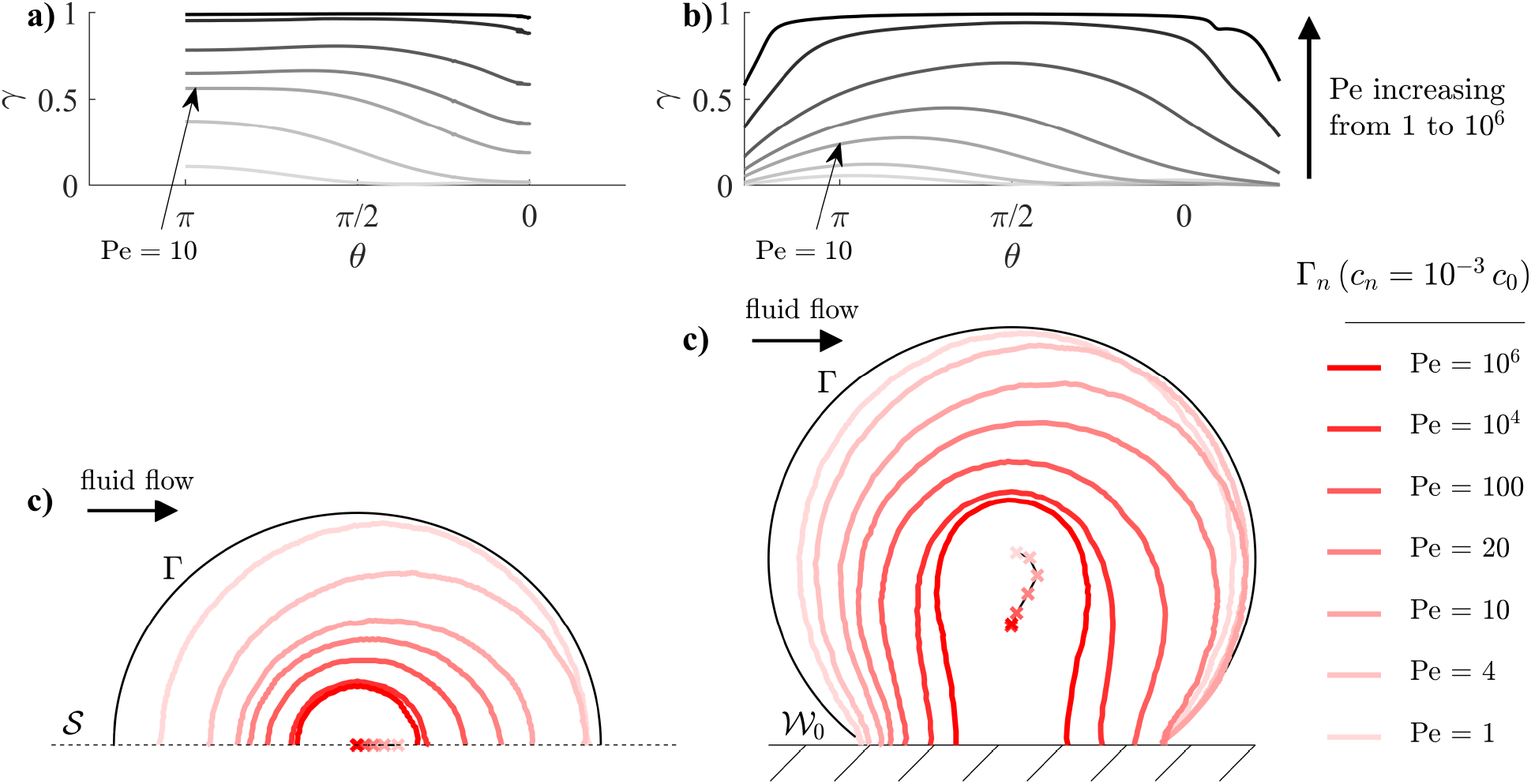
Solute concentration in microfluidic culture for Da = 5, R_d_ = 0.5 (above the onset of rate-induced necrosis). **a)**. surface concentration, unconfined geometry. **b)**. id., confined geometry. **c)**. necrotic boundaries for varying Pe, unconfined geometry. **d)**. id., confined geometry.

To estimate the configuration of maximum asymmetry of solute concentration, we consider the quasi-1D domain 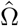 and derive the conditions where 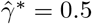 (Section SR6). Those are

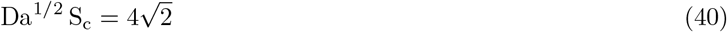

The quantity Da^1/2^ S_c_ is a good predictor of the maximum asymmetry of simulated necrotic cores (Figure 11). Using dimensional parameters,

**Figure 11.**
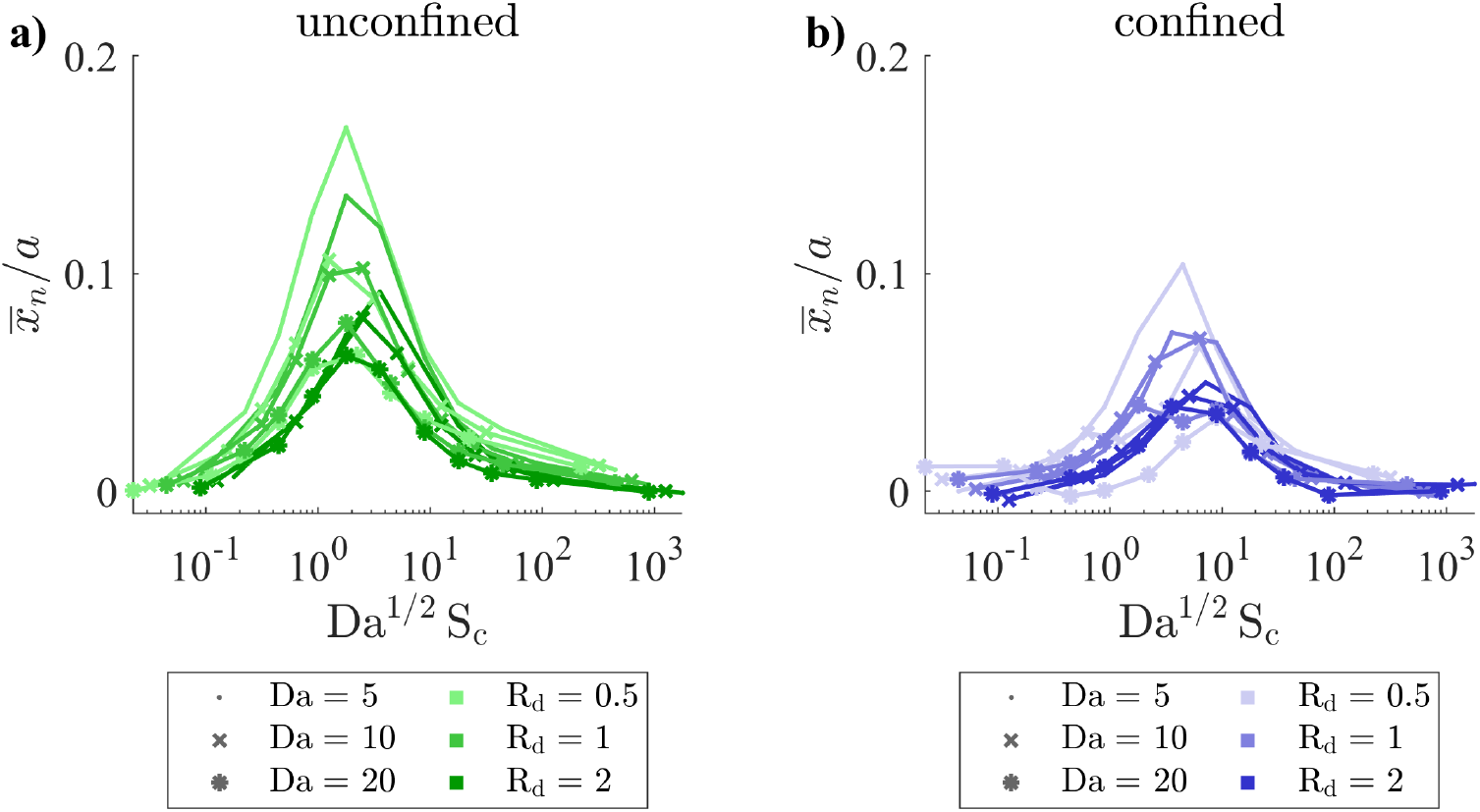
Trajectory of necrotic barycentres in microfluidic culture (*x*-coordinate). **a)**. unconfined geometry. **b)**. confined geometry.

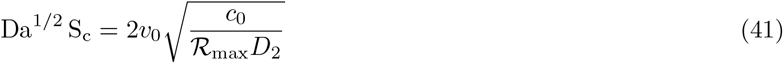

The symmetric nature of solute concentration thus only depends on solute kinetics inside M-CELS and on boundary conditions on ℐ. Since it does not depend on the M-CELS diameter or on the microfluidic channel dimensions, the peak inlet velocity could be used as a single parameter to induce persistent asymmetric concentrations inside M-CELS and investigate directional responses. The maximum downstream shift of the simulated necrotic barycentres occurrs for Da^1/2^ S_c_ ∈ [1, 4] in the unconfined geometry (Figure 11a). The shift was delayed and less marked in the confined geometry because solute supply increases more slowly near the recirculation areas. It occurred at Da^1/2^ S_c_ ∈ [3, 10] (Figure 11b).

#### 3.3.5 Approach of maximal-supply asymptote

Let us consider cases of rate-induced necrosis (Da > Da_∞_) and assume that convection is sufficiently high that the asymmetry peak has passed i.e. , Da^1/2^ S_c_ > 4 in the unconfined geometry and Da^1/2^ S_c_ > 10 in the confined geometry. The approach of the maximal-supply asymptote depends on the rate at which solute supply increases on Γ via the diffusion-dominated areas. Let us consider the recirculation areas and define a corner Péclet number Pe_*ρ*_ based on the recirculation properties and the quasi-1D no-flow solution:

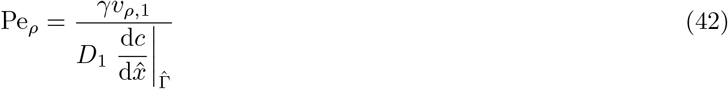

*v*_*ρ*,1_ is the maximum velocity of the largest eddy, with *v*_*ρ*,1_ = *qv*_0_ and *q* ∼ 10^−3^ (Fig. 5 in [28]). From the no-flow expression of concentration in 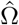, obtained by substituting (70) into (68), we obtain

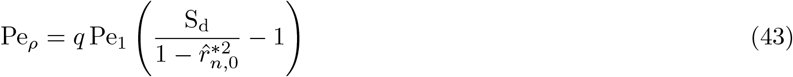

Substituting 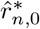 with 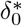 using Equation (74) and expanding at order 1 in 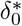 gives

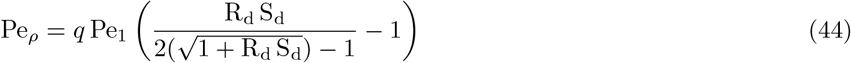

Under deficient diffusive supply (R_d_ S_d_ ⪡ 1), successive Taylor expansions lead to

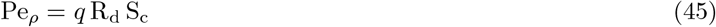

The balance between convective supply and consumption on Γ_*ρ*_ is defined, analogously to S_c_, as

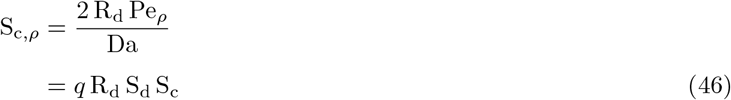

A similar reasoning may be followed for the diffusive boundary layer covering all of Γ. The velocity of the first eddy *qv*_0_ would be replaced by the velocity at the entrance of the boundary layer, which also scales with *v*_0_. Solute supply to Γ thus is governed by R_d_ S_d_ S_c_ when S_c_ ⪢ 1. The excess of convective supply required to maintain a given proportion of live cells (Φ_*L*_) scales with (R_d_ S_d_)^−1^. The corresponding peak inlet fluid velocity scales like

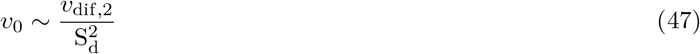

The approach of maximal supply via external convection thus is delayed by deficient diffusive supply to the M-CELS and by fast diffusion within it. Using dimensional parameters,

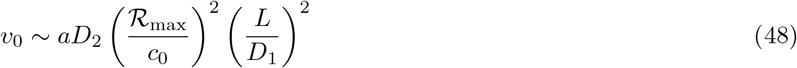

This velocity evolves linearly with the M-CELS diameter, so it would need to increase with M-CELS age if solute penetration distance were to be maintained over the course of an experiment. In the example of neurons in a cortical organoid, the increase in radial diffusivity resulting from neuronal alignment would also contribute to *v*_0_ needing to increase with M-CELS age. The dependency of *v*_0_ on the square of the inverse of the inlet concentration and on the square of the distance to the solute source provide means to reduce experimentally-required flow rates. These come with the caveats that inlet concentrations often are physically or biologically constrained (e.g. , dissolved gases like oxygen have a saturation limit) and that the distance to the solute source is constrained.

The numerical simulations verified that the rate of approach of maximal supply is governed by R_d_ S_d_ S_c_. In the unconfined geometry, overcoming the diffusive boundary layer on Γ to achieve 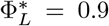 required R_d_ S_d_ S_c_ > 10 (Figure 12a). In the confined geometry, the additional diffusive barrier constituted by the corner recirculation areas made that condition R_d_ S_d_ S_c_ > 10^3^. This limits the biological configurations where supply deficiency can be filled within the constraints of shear stress on cells.

**Figure 12.**
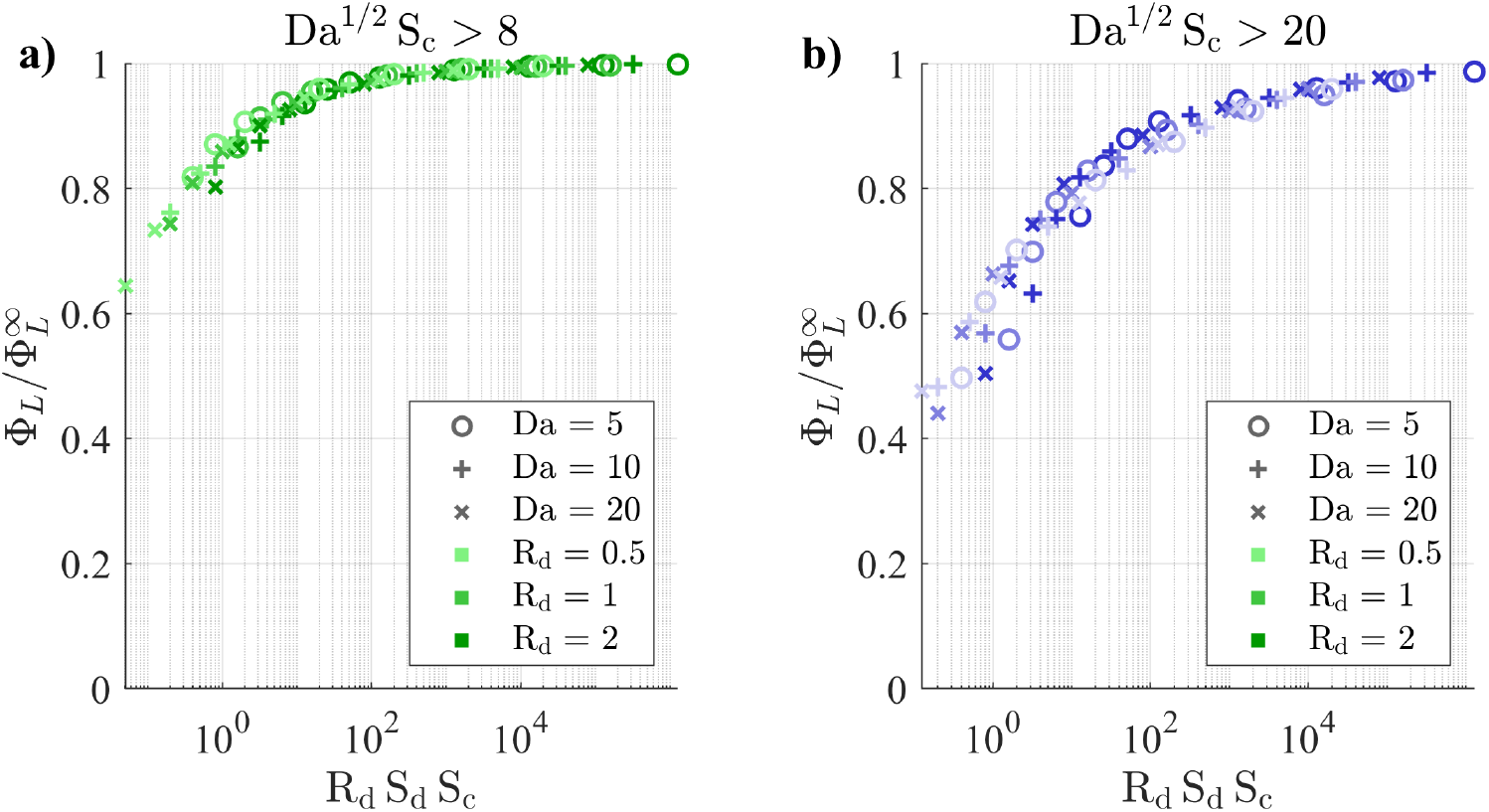
Approach of maximal supply via increased convective solute supply to Γ. The live fraction is governed by R_d_ S_d_ S_c_. **a)**. unconfined geometry. **b)**. confined geometry

## 4 Discussion

Multi-cellular engineered living systems (M-CELS) are situated at a certain distance from the source of soluble nutrients, growth factors, or drugs, that need to be supplied to them. Since solute transport is diffusion-dominated within M-CELS, internal solute distribution is linked to external solute transport via solute concentration at the fluid–M-CELS interface only. It thus is key to rapidly cross the source–M-CELS distance. Both slow external transport relative to M-CELS metabolic needs and a large surface area of contact with the culture-chamber walls limit solute distribution in M-CELS.

We quantified the maximum consumption-to-diffusion kinetics ratio, noted Da_∞_, that an M-CELS can reach without necrosis as a function of its structural confinement. Finite transport kinetics in microfluidic chambers imply the need for an excess of solute supply to the M-CELS surface. That excess is a hyperbolic function of the ratio Da / Da_∞_. The cost of preventing necrosis thus diverges when Da approaches Da_∞_ e.g., requiring infinite flow rates. This renders the realisation of a non-necrotic M-CELS at Da_∞_ unfeasible and places a lower limit on the maximum consumption-to-diffusion ratio that can be achieved in a given experimental setup. That limit depends on design constraints of the setup and on cellular sensitivity to shear stress.

External convection brings about a higher solute supply to the fluid–M-CELS interface only after the solutes have crossed zones where their transport is diffusion-dominated. These cover the whole fluid–M-CELS interface and include the diffusive boundary layer that grows from the interface into the fluidic chamber and the recirculation area near each confinement corner. The latter exists at the intersection of the M-CELS and the wall in the cylindrical geometry we simulated. It would be present near any corner of angle inferior to 146° in the confinement structure of a spherical M-CELS [28]. Slow diffusion across those areas relative to solute consumption in the M-CELS impedes the benefits of external convection. This arises when the diffusion-dominated areas are thick e.g., when there are narrow corners, and when diffusive supply in an equivalent static configuration is deficient e.g., when the distance between solute source and M-CELS is large. In both cases, slow diffusion kinetics lower concentration on the fluid–M-CELS interface and external convection needs to be all the higher to overcome the diffusive barrier. The same reasoning applies to M-CELS encapsulated in hydrogels. Solute diffusivity across hydrogels is close to their diffusivity in water thanks to high hydrogel porosity, but capsule thickness is detrimental to increased solute penetration via external convection. One should thus aim for capsules to be as thin as possible so as not to impede solute supply whilst maintaining the cell–matrix interactions and the protection against shear stress that these provide.

The easiest means for experimentalists to increase solute supply to fluid–M-CELS interfaces is to modulate transport velocities, either by design for diffusion (short distance between solute source and M-CELS) or by operation for convection (peak fluid velocity). The effect of both is penalised by fast diffusion within the M-CELS, independently of how the internal-diffusion kinetics compare with consumption and external-diffusion kinetics. Fast internal diffusion means that any increase in solute supply to the fluid–M-CELS interface is more rapidly distributed throughout the M-CELS, slowing down the increase in interface concentration that is necessary to increase solute penetration depth. Achieving non-necrotic M-CELS (or a given penetration depth) at a given Da thus requires less external convection in large M-CELS with low consumption than in small M-CELS with high consumption.

The fundamental understanding of how microfluidic chamber design and operation affect solute supply to M-CELS that we have presented allows to state how much larger rigid avascular M-CELS can grow by adding fluid flow to the culture setup. In the oxygen example that illustrated our results, the maximum no-flow Da was a fourth of Da_∞_ i.e., the maximum radius was divided by 2. Fluid flow around an unconfined M-CELS allowed to recover a maximal Da close to Da_∞_ (Figure 9a). Peak fluid velocity is constrained in experimental setups because of the shear-stress sensitivity of cells, which capped the maximum achievable Da at Da_∞_ /2 in the presence of recirculation areas i.e., an increase in maximal radius of around 40% compared to static culture (Figure 9b). Similarly, the approach of maximal supply in an M-CELS with incomplete solute penetration required a 100-times faster fluid flow because of the need for solutes to cross the recirculation areas (Figure 12b). This highlights the importance of eliminating thick diffusion-dominated areas in confinement structures. These exist in widely used U-shaped barriers and wells (e.g., [30]), but are absent from designs such as slit U-shaped wells with smooth contours [31] or porous confinement scaffolds [32]. These result also point to the inhibition of solute-transport benefits by the encapsulation of M-CELS in hydrogels.

Current experimental efforts in organoid research focus on incorporating a perfusable vasculature to better mimick *in vivo* mechanical forces and solute-transport patterns. This study of the link between the geometry and operation of the culture setup and solute concentration inside M-CELS can inform the boundary conditions at the entrance of the vessels. This would in turn contribute to quantifying the inhomogeneity of solute transport in perfusable M-CELS as well.

## Data availability

The simulation results and processing codes are on the GitHub repository solute_transport_mcels_microfluidic.

## Acknowledgements

All authors gratefully acknowledge support from T. Christian Gasser. WB and SB gratefully acknowledge support from the Department of Engineering Mechanics, School of Engineering Sciences, KTH. This project has received funding from the Olle Engkvist foundation (OES 213-0231), the Knut and Alice Wallenberg Foundation (KAW 2021.0172) and the European Research Council under the European Union’s Horizon Europe research and innovation programme (PHOENIX grant agreement 101043985) via grants led by MT.

