## Supplementary Information for "Capacity and limitations of microfluidic flow to increase solute transport in three-dimensional cell cultures"

#### Supplementary Methods

##### SM1 Quasi-1D analytical domain

In order to acquire an analytical understanding of the problem, we define a quasi-1D "radial-linear" domain $\hat{\Omega}$ as the union of a radial section of a disc of radius $a$ and angle $\mathrm{d}\theta$, named $\hat{\Omega}_2$ and centered on $\hat{O}$, and the constant-width extension of that section, named $\hat{\Omega}_1$ (Supplementary Figure S1). Let $\hat{\Gamma} = \hat{\Omega}_1 \cap \hat{\Omega}_2$ and $L$ be the distance between $\hat{\Gamma} \cap \{\hat{y} = 0\}$ and the inlet plane of $\hat{\Omega}_1$, named $\hat{\mathcal{I}}$. $\hat{\Omega}$ is described by the cartesian coordinate system $(\hat{O}, \hat{x}, \hat{y})$ and the associated polar coordinate system $(\hat{O}, \hat{r}, \hat{\theta})$. The planes $\hat{\mathcal{S}}_r = \{\hat{\theta} = \pm\,\mathrm{d}\theta/2\}$ are assumed to be planes of symmetry for $\hat{\Omega}_2$ and the planes $\hat{\mathcal{S}}_l$ of normal $\hat{y}$ that bound $\hat{\Omega}_1$ are assumed to be planes of symmetry for $\hat{\Omega}_1$. If $\mathrm{d}\theta \ll 1$, we assume that, for a scalar $\xi$

$$\left.\frac{\mathrm{d}\xi}{\mathrm{d}\hat{\mathbf{r}}} \cdot \hat{\mathbf{r}}\right|_{\hat{\Gamma}} = \left.\frac{\mathrm{d}\xi}{\mathrm{d}\hat{\mathbf{x}}} \cdot \hat{\mathbf{x}}\right|_{\hat{\Gamma}} \tag{49}$$

The assumptions of symmetry and narrowness of $\hat{\Omega}_2$ allow equations in $\hat{\Omega}$ to be reduced to a quasi-1D "radial-linear" form where variables depend on $\hat{x}$ only in $\hat{\Omega}_1$ and on $\hat{r}$ only in $\hat{\Omega}_2$.

##### SM2 Fluid flow equations

Fluid flow in $\Omega_1$ was modelled by the steady Navier-Stokes equations with a parabolic inlet velocity and normal outlet flow:

$$\rho(\mathbf{v}_1 \cdot \boldsymbol{\nabla})\mathbf{v}_1 = -\boldsymbol{\nabla} p + \mu \boldsymbol{\nabla} \cdot (\boldsymbol{\nabla}\mathbf{v}_1 + \boldsymbol{\nabla}\mathbf{v}_1^T) \tag{50a}$$

$$\boldsymbol{\nabla} \cdot \mathbf{v}_1 = 0 \tag{50b}$$

$$\mathbf{v}_1|_{\mathcal{I}} = v_0 \left(1 - \left(\frac{y}{h}\right)^2\right) \mathbf{e}_x \tag{50c}$$

$$p_1|_{\mathcal{O}} = 0 \tag{50d}$$

$$\mathbf{v}_1 \cdot \mathbf{e}_y|_{\mathcal{O}} = 0 \tag{50e}$$

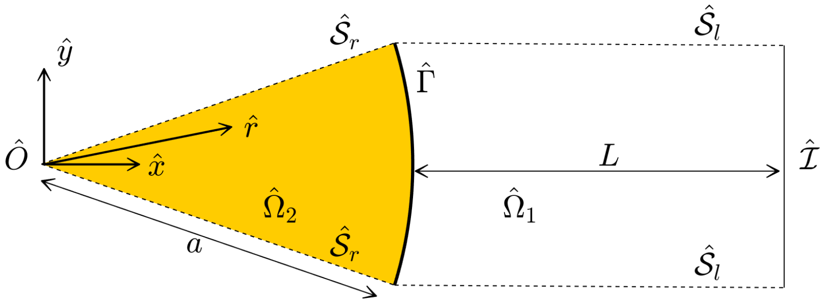


Figure S1: Quasi-1D "radial-linear" reduction of $\Omega$ into $\hat{\Omega}$. The angular section $\hat{\Omega}_2$ has an angle $\mathrm{d}\theta \ll 1$.

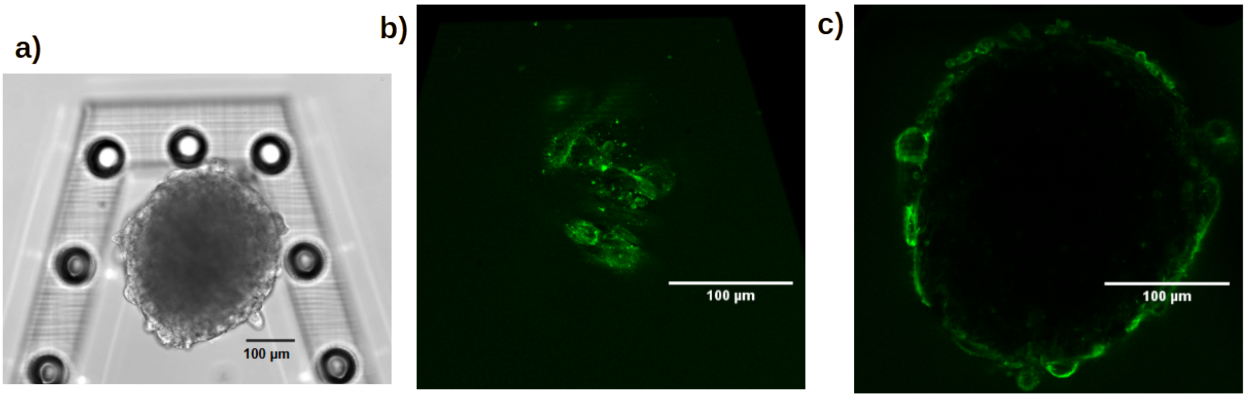


Figure S2: Human lung-fibroblast spheroid (3000 seeded cells, day 3). **a)**. Brightfield view of spheroid loaded in microfluidic chamber between confinement pillars. **b)**. GFP staining of the cytoskeletal protein actin, 0.1 µm above the bottom of the chamber. **c)**. id., 55 µm above the bottom of the chamber.

We assume that the imposed time scales of fluid flow are larger than the convective time scale and therefore neglect local acceleration. Let non-dimensionalised variables be defined as $\mathbf{x} = \mathbf{x}^* a$, $\mathbf{v}_1 = \mathbf{v}_1^* v_0$, and $p = p^* p_0$, where $p_0$ is such that the pressure drop along $\Omega_1$ is of the order of $p_0$. Parabolic flow at the inlet dictates that $v_0 \sim Gh^2/\mu$, where $G$ is the pressure gradient driving the flow. Here, because of the constriction between the apex of $\Omega_2$ and the wall at $y = 2h$, most of the pressure drop occurs when fluid flows over the M-CELS, which means $G \sim p_0/a$. Since $a \sim h$, $v_0 \sim p_0/(a\mu)$. The normalised momentum conservation equation (50a) becomes, dropping the asterisks for concision:

$$\mathrm{Re}(\mathbf{v}_1 \cdot \boldsymbol{\nabla})\mathbf{v}_1 = -\boldsymbol{\nabla} p + \boldsymbol{\nabla} \cdot (\boldsymbol{\nabla}\mathbf{v}_1 + \boldsymbol{\nabla}\mathbf{v}_1^T) \tag{51}$$

where the Reynolds number has been defined as $\mathrm{Re} = \rho v_0 a/\mu$.

#### SM3 Supplementary numerical methods

The packages *Laminar Flow*, *Darcy's law*, and *Transport of Diluted Species* of COMSOL Multiphysics were used. In the *Laminar flow* package, the discretisation was changed to P2+P1. In order to avoid sharp corners, the geometry at the contact points $\Omega_1 \cap \Omega_2 \cap \mathcal{W}$ was modified by inserting short edges of normal $\mathbf{x}$ and length 10 µm between $\Gamma$ and $\mathcal{W}$. The mesh applied to $\Omega$ was triangular. Its size independence was evaluated on the normal shear rate $(\mathrm{d}\mathbf{v}_1/\mathrm{d}\mathbf{n}) \cdot \mathbf{n}$ and the normal concentration gradient $(\mathrm{d}c_1/\mathrm{d}\mathbf{n}) \cdot \mathbf{n}$ on $\Gamma$ and on the concentration $c_2$ on the line of normal $y$ passing through the center of $\Omega_2$. The grid was refined by dividing the number of elements by 2 until each of the criteria changed by less than 1% between two successive refinement steps. This yielded a triangular mesh of maximum size 25 µm in $\Omega$. The interface $\Gamma$ was meshed at 6 µm and a boundary layer was grown from it into $\Omega_1$ and $\Omega_2$ with a

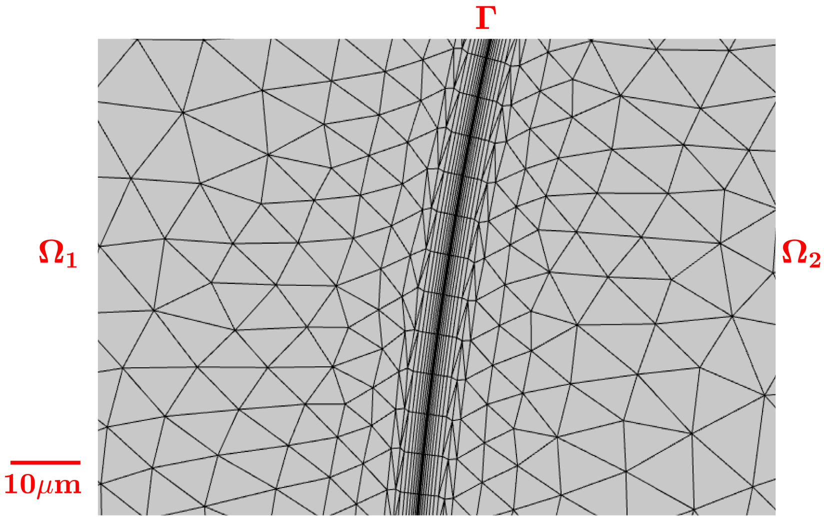


Figure S3: Mesh used in the numerical simulations (zoom onto the fluid–M-CELS interface

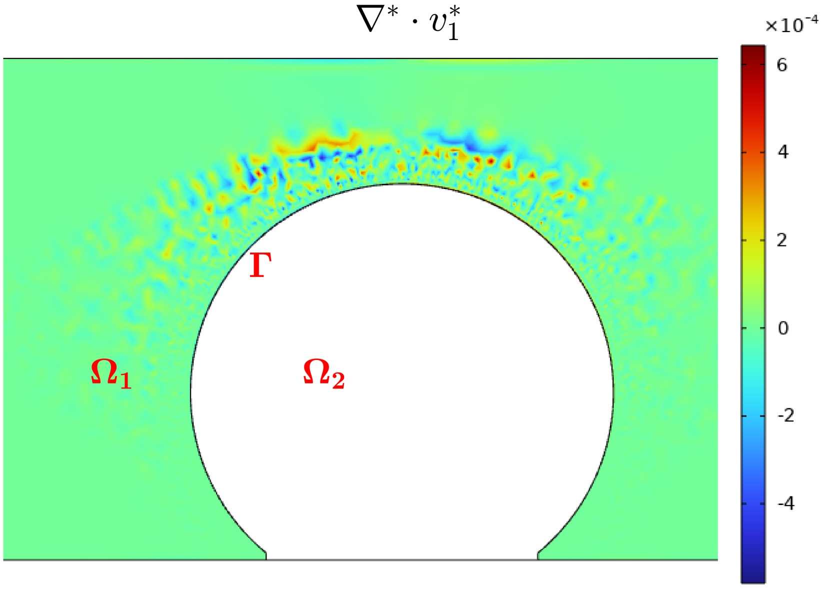


Figure S4: Divergence of fluid velocity in $\Omega_1$. $v_0 = 5$ mm/s in the displayed example.

first-layer thickness of 2 µm and a stretching factor of 1.2 over 8 layers (Supplementary Figure S3). The laminar-flow solution was accurate to $|\boldsymbol{\nabla}^* \cdot \mathbf{v}_1^*| < 10^{-3}$ (Supplementary Figure S4).

Simulations at the static asymptote were conducted by sweeping the following lattice:

$$\mathrm{Da} \in \{1, 2, 3, 5, 10, 20\}$$

$$\mathrm{R_d} \in \{0.1, 0.25, 0.5, 1, 2, 4, 10, 20, 100\}$$

This was implemented by varying $\mathcal{R}_{\mathrm{max}}$ and $D_2$, with the other parameters fixed as per Table S1.

Fluidic simulations were conducted by sweeping the following lattice for necrosis prevention:

$$\mathrm{Da} \in \{1, 2, 3\}$$

$$\mathrm{R_d} \in \{0.25, 0.5, 1\}$$

Table S1: Fixed simulation parameters.

| Name | Symbol | Value |
|---|---|---|
| fluid-channel length | $L$ | 6 mm |
| fluid-channel height | $h$ | 0.8 mm/1.2 mm(1) |
| M-CELS radius | $a$ | 0.5 mm |
| inlet solute concentration | $c_0$ | 0.2 mol/m$^3$ |
| half-rate constant | $K_{1/2}$ | 4.6 10$^{-5}$ mol/m$^3$ |
| diffusivity in fluid | $D_1$ | 1e-9 m$^2$/s(2) |

(1) unconfined/confined
(2) for $\mathrm{Pe}_1 \leq 10^4$. Multiplied by $10^4/\mathrm{Pe}_1$ for $\mathrm{Pe}_1 > 10^4$ to keep $\mathrm{Re} < 1$

Table S1: Fixed simulation parameters.

| Name | Symbol | Value |
|---|---|---|
| distance inlet–M-CELS | $L$ | 2.5 mm |
| fluid-channel height | $h$ | 0.8 mm/1.2 mm(1) |
| M-CELS radius | $a$ | 0.5 mm |
| inlet solute concentration | $c_0$ | 0.2 mol/m$^3$ |
| half-rate constant | $K_{1/2}$ | 4.6 10$^{-5}$ mol/m$^3$ |
| diffusivity in fluid | $D_1$ | 1e-9 m$^2$/s(2) |

(1) unconfined/confined
(2) for $\mathrm{Pe}_1 \leq 10^4$. Multiplied by $10^4/\mathrm{Pe}_1$ for $\mathrm{Pe}_1 > 10^4$ to keep $\mathrm{Re} < 1$

$$\mathrm{Pe}_1 \in \{10^{-1}, 0.25, 0.5, 1, 2, 4, 10, 20, 40, 10^2\}$$

and the following lattice for the approach of the maximal-supply asymptote:

$$\mathrm{Da} \in \{5, 10, 20\}$$
$$\mathrm{R_d} \in \{0.5, 1, 2\}$$
$$\mathrm{Pe}_1 \in \{10^{-1}, 0.25, 0.5, 1, 2, 4, 10, 20, 40, 10^2, 10^3, 10^4, 10^5, 10^6\}$$

For $\mathrm{Pe}_1 \leq 10^4$, this was done by varying $v_0$. In order to keep $\mathrm{Re} < 1$, simulations at $\mathrm{Pe}_1 \geq 10^5$ were done at the velocity set for $\mathrm{Pe}_1 = 10^4$ with diffusivities and consumption rate divided by 10 for each increasing power of 10 of $\mathrm{Pe}_1$.

### Supplementary Results

#### SR1 Analytical expression of the necrotic radius at the maximal-supply asymptote

Solute concentration is radially symmetric at the maximal-supply asymptote. The transport equation (8) becomes, with the approximation of a constant consumption rate:

$$0 = \frac{1}{r^*}\frac{\mathrm{d}}{\mathrm{d}r^*}\left(r^*\frac{\mathrm{d}c_2^*}{\mathrm{d}r^*}\right) - \mathrm{Da}\,\mathbf{1}_{c_2\geq 0} \tag{52}$$

where $\mathbf{1}_I$ is the indicator function of the space where the condition $I$ is verified. It is subjected to the maximal-supply boundary condition

$$c_2^*|_{r^*=1} = 1 \tag{53}$$

We introduce the necrotic radius $r_n$ such that $\{c_2 = 0\} = \{r \leq r_n\}$. In the unapproximated problem with Michaelis-Menten kinetics, the $\mathcal{C}^1$-regularity of $c_2$ implies that $c_2$ is zero with a zero gradient at $r_n$. We thus assume that $r_n$ is such that

$$c_2|_{r=r_n} = 0 \tag{54a}$$

$$\left.\frac{\mathrm{d}c_2}{\mathrm{d}r}\right|_{r=r_n} = 0 \tag{54b}$$

The integration of Equation (52) between $r_n^*$ and 1, subject to (54a) and (54b), gives

$$c_2^* = \frac{\mathrm{Da}}{4}\left(r^{*2} - r_n^{*2}\right) - \frac{\mathrm{Da}}{2}r_n^{*2}\ln\left(\frac{r}{r_n}\right) \tag{55}$$

with $r_n$ such that (53) is satisfied i.e. , it verifies

$$1 = \frac{\mathrm{Da}}{4}\left(1 - r_n^{*2}\right) + \frac{\mathrm{Da}}{2}r_n^{*2}\ln(r_n^*) \tag{56}$$

If the necrotic area is large, $r_n^* \approx 1$ and $\mathrm{Da} \gg 1$. Let $\delta^* = 1 - r_n^*$. The logarithm term may be expanded around $\delta^* = 0$ as

$$\begin{aligned}\ln\left(r_n^*\right) &= \ln\left(1 - \delta^*\right)\\ &= -\delta^* - \frac{\delta^{*2}}{2}\end{aligned}$$

which yields the following expansion of Equation (56) at order 2 in $\delta^*$:

$$\frac{\mathrm{Da}}{2}\delta^{*2} - 1 = 0 \tag{57}$$

The asymptotic expressions for $r_n^*$ and $\Phi_L^\infty$ at high Da follow:

$$r_n^* = 1 - \left(\frac{2}{\mathrm{Da}}\right)^{1/2} \tag{58}$$

$$\Phi_L^{\infty} = 2\left(-\frac{1}{\mathrm{Da}} + \left(\frac{2}{\mathrm{Da}}\right)^{1/2}\right) \tag{59}$$

#### SR2 Analytical expression of the necrotic radius at the static asymptote

In $\hat{\Omega}$, the boundary value problem approximated with a constant consumption rate admits the following (dimensional) equations:

$$0 = D_1 \frac{\mathrm{d}^2 c_1}{\mathrm{d}\hat{x}^2}, \quad \hat{x} \in \hat{\Omega}_1 \tag{60}$$

$$0 = \frac{D_2}{\hat{r}} \frac{\mathrm{d}}{\mathrm{d}\hat{r}}\left(\hat{r}\frac{\mathrm{d}c_2}{\mathrm{d}\hat{r}}\right) - \mathcal{R}_{\max}, \quad \hat{r} \in \hat{\Omega}_2 \tag{61}$$

We assume that a necrotic core exists and is defined by the equiradial surface $\hat{r} = \hat{r}_n$. The boundary conditions are

$$c_1|_{\hat{x}=a+L} = c_0 \tag{62}$$

$$c_2|_{\hat{r}=\hat{r}_n} = 0 \tag{63}$$

$$\left.\frac{\mathrm{d}c_2}{\mathrm{d}\hat{r}}\right|_{\hat{r}=\hat{r}_n} = 0 \tag{64}$$

(65)

The continuity conditions on $\hat{\Gamma}$ are

$$c_1|_{\hat{x}=a^+} = c_2|_{\hat{r}=a^-} \tag{66}$$

$$D_1 \left.\frac{\mathrm{d}c_1}{\mathrm{d}\hat{x}}\right|_{\hat{x}=a^+} = D_2 \left.\frac{\mathrm{d}c_2}{\mathrm{d}\hat{r}}\right|_{\hat{r}=a^-} \tag{67}$$

where $\hat{x} = a^+$ (respectively $\hat{r} = a^-$) signifies $\{\hat{x} \to a, \hat{x} > a\}$. The integration of (60) between $a$ and $a + L$ gives

$$c_1 = A_1(\hat{x} - (L + a)) + c_0, \quad \hat{x} \geq a \tag{68}$$

The integration of (61) between $\hat{r}_n$ and $a$ subject to the zero-flux condition at $\hat{r}_n$ (64) gives:

$$c_2 = \frac{\mathcal{R}_{\max}}{4D_2}(\hat{r}^2 - \hat{r}_n^2) - \frac{\mathcal{R}_{\max}}{2D_2}\hat{r}_n^2 \ln\left(\frac{\hat{r}}{\hat{r}_n}\right), \quad \hat{r} \leq a \tag{69}$$

The continuity of diffusive flux on $\Gamma$ (67) implies

$$A_1 = \frac{\mathcal{R}_{\max} a}{2D_1}\left(1 - \left(\frac{\hat{r}_n}{a}\right)^2\right) \tag{70}$$

which yields the following equation for $\hat{r}_n^* = \hat{r}_n/a$ after subsitution into the concentration continuity condition (66):

$$-\frac{1}{\mathrm{S_d}}(1 - \hat{r}_n^{*2}) + 1 = \frac{\mathrm{Da}}{4}(1 - \hat{r}_n^{*2}) + \frac{\mathrm{Da}}{2}\hat{r}_n^{*2} \ln(\hat{r}_n^*) \tag{71}$$

with the definitions of Da and $\mathrm{S_d}$ given in Sections 2.3 and 2.4.

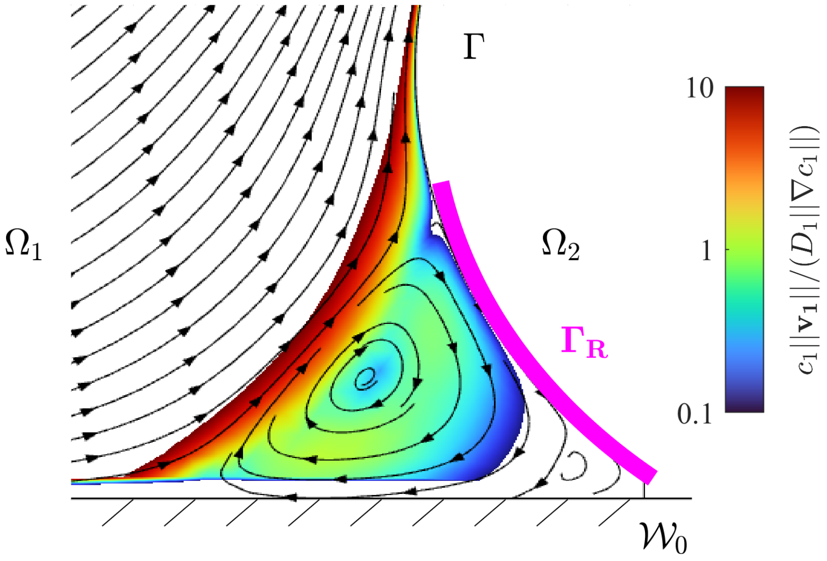


Figure S5: Streamlines of $\mathbf{v}_1$ in the vicinity of $\Gamma \cap \mathcal{W}_0$ with the first and second Moffatt eddies visible near the corner. $\Gamma_\rho$ defined as the portion of $\Gamma$ adjacent to the recirculation eddies. Coloured surface: local ratio of magnitudes of convective to diffusive fluxes for $Pe_1 = 10^4$. Ratios are shown when they are between 0.1 and 10. They are under 0.1 in the blank region adjacent to $\Gamma \cap \mathcal{W}_0$ and above 10 in the other blank regions of $\Omega_1$.

#### SR3 Approach of the maximal-supply asympote in static culture

We now investigate the expression of the live fraction as a function of diffusive supply when there is rate-induced insufficient transport i.e. , $\delta_0^* = 1 - \hat{r}_{n,0}^* \ll 1$. In this asymptotic expansion, we assume that diffusive supply is ample i.e. , $\mathrm{S_d} \gg 1$, and $\mathrm{R_d} \gg 1$. The same Taylor expansion of the logarithm term as for the maximal-supply asymptote transforms (31) into

$$\left(-\frac{1}{\mathrm{S_d}} + \frac{\mathrm{Da}}{2}\right)\delta_0^{*2} + \frac{2}{\mathrm{S_d}}\delta_0^* - 1 = 0 \tag{72}$$

which $\mathrm{R_d} \gg 1$ allows to approximate as

$$\delta_0^{*2} + \frac{2}{\mathrm{R_d}}\delta_0^* - \frac{2}{\mathrm{Da}} = 0 \tag{73}$$

This admits a positive solution

$$\delta_0^* = -\frac{1}{\mathrm{R_d}}\left(1 - \sqrt{1 + \mathrm{R_d}\,\mathrm{S_d}}\right) \tag{74}$$

The live fraction normalised by its value at the maximal-supply asymptote is then, at order 1 in $\delta^*$:

$$\begin{aligned}\frac{\Phi_L^0}{\Phi_L^\infty} &= \frac{\pi(a^2 - \hat{r}_{n,0}^{*2})}{\pi(a^2 - \hat{r}_{n,\infty}^{*2})} \\ &= \frac{-\delta_0^{*2} + 2\delta_0^*}{-\delta_\infty^{*2} + 2\delta_\infty^*} \\ &\approx \frac{\delta_0}{\delta_\infty}\end{aligned}$$

which may be expressed as

$$\frac{\Phi_L^0}{\Phi_L^\infty} = -(\mathrm{R_d}\,\mathrm{S_d})^{-1/2} + \left(1 + (\mathrm{R_d}\,\mathrm{S_d})^{-1}\right)^{1/2} \tag{75}$$

#### SR4 Streamlines near the confinement corner

#### SR5 Analytical expression of the necrotic radius in microfluidic culture

Let us assume an M-CELS with $\mathrm{Da} < \mathrm{Da}_\infty$ in the radial-linear domain $\hat{\Omega}$. The quality of the approximation made by using $\hat{\Omega}$ is reduced in presence of fluid flow because solute transport is not symmetric with respect to $\{x = 0\}$, unless $\mathrm{S_c} \gg \mathrm{Da}$. In that case, the rate of consumption is small compared to the rate at which solute flows over the M-CELS and convected solutes reach the downstream side of the M-CELS at an unchanged concentration. We assume that this condition is necessary for the prevention of necrosis, so that concentration in a real system is at least approximately symmetric.

Let convection be uniform at $v_0$ in $\hat{\Omega}_1$ and let there be a necrotic radius $\hat{r}_n$ such that $0 \leq \hat{r}_n < a$. The transport equation in $\hat{\Omega}_1$ is

$$0 = D_1 \frac{\mathrm{d}^2 c_1}{\mathrm{d}\hat{x}^2}, \quad \hat{x} \in \hat{\Omega}_1 \tag{76}$$

Equation (61) holds in $\hat{\Omega}_2$, like in the static asymptote. Application of the Dirichlet condition on $\mathcal{I}$ and of the diffusive-flux continuity condition on $\hat{\Gamma}$ yields the following expression for the concentration in $\hat{\Omega}_1$:

$$c_1 = c_0 + \frac{\mathcal{R}_{\max}}{2v_0}\left(a - \frac{\hat{r}_n^2}{a}\right)\left(e^{-\mathrm{Pe}_1} - e^{-\dfrac{v_0\hat{x}}{D_1}}\right) \tag{77}$$

The concentration in $\hat{\Omega}_2$ satisfies Equation (69). Applying concentration continuity on $\hat{\Gamma}$ yields the following non-dimensionalised equation for $\hat{r}_n^*$:

$$\frac{e^{-\mathrm{Pe}_1} - 1}{\mathrm{S_c}}(1 - \hat{r}_n^{*2}) + 1 = \frac{\mathrm{Da}}{4}(1 - \hat{r}_n^{*2}) + \frac{\mathrm{Da}}{2}\hat{r}_n^{*2}\ln(\hat{r}_n^*) \tag{78}$$

#### SR6 Conditions of maximum necrotic-core asymmetry in microfluidic culture

In order to estimate the configuration of maximum asymmetry of solute concentration, we consider the analytical domain $\hat{\Omega}$ and derive the conditions where $\hat{\gamma}^* = 0.5$. Equation (77) specialised at $x = a$ yields

$$1 - \frac{1}{\mathrm{S_c}}(1 - r_n^{*2}) = \frac{1}{2} \tag{79}$$

Assuming that necrotic cores are still large at this stage and expanding at the first order in $\delta^* = 1 - r_n^*$, this equation becomes

$$1 - \frac{2\delta^*}{\mathrm{S_c}} = \frac{1}{2} \tag{80}$$

The concentration in $\hat{\Omega}_2$ obeys (69), leading to

$$\left(-\frac{1}{\mathrm{S_c}} + \frac{\mathrm{Da}}{2}\right)\delta^{*2} + \frac{2}{\mathrm{S_c}}\delta^* - 1 = 0 \tag{81}$$

Substituting the positive solution of the latter expression into (80) gives

$$1 - \frac{2}{\dfrac{\mathrm{Da}\,{\mathrm{S_c}}^2}{2} - \mathrm{S_c}} \left( 1 + \left( 1 + \frac{\mathrm{Da}\,{\mathrm{S_c}}^2}{2} - \mathrm{S_c} \right)^{1/2} \right) = \frac{1}{2} \tag{82}$$

The maximum asymmetry of solute supply to $\Gamma$ corresponds to a case where supply is abundant on the upstream side of $\Omega_2$ but deficient on the downstream side. We thus may assume $\mathrm{S_c} > 1$. The inequality $\mathrm{Da}\,{\mathrm{S_c}}^2 \gg \mathrm{S_c}$ is verified in all cases of rate-induced necrosis ($\mathrm{Da} > 4$) and the above expression may be simplified into

$$\mathrm{Da}^{1/2}\,\mathrm{S_c} = 4\sqrt{2} \tag{83}$$
